# SQUARNA - an RNA secondary structure prediction method based on a greedy stem formation model

**DOI:** 10.1101/2023.08.28.555103

**Authors:** Davyd R. Bohdan, Grigory I. Nikolaev, Janusz M. Bujnicki, Eugene F. Baulin

## Abstract

Non-coding RNAs play a diverse range of roles in various cellular processes, with their spatial structure being pivotal to their function. The RNA’s secondary structure is a key determinant of its overall fold. Given the scarcity of experimentally determined RNA 3D structures, understanding the secondary structure is vital for discerning the molecule’s function. Currently, there is no universally effective solution for de novo RNA secondary structure prediction. Existing methods are becoming increasingly complex without marked improvements in accuracy, and they often overlook critical elements such as pseudoknots. In this work, we introduce SQUARNA, a novel approach to de novo RNA secondary structure prediction. This method utilizes a simple, greedy stem formation model, addressing many of the limitations inherent in previous tools. Our benchmarks demonstrate that SQUARNA matches the performance of leading methods for single sequence inputs and significantly surpasses existing tools when applied to sequence alignment inputs.

## INTRODUCTION

Hundreds of structured RNA molecules have been identified and characterized to date [1], ranging from ligand-binding riboswitches that regulate gene expression [2] to structured elements in viral RNAs that confer resistance to degradation [3]. The function of these RNAs is governed by their spatial structure [4] which in turn is determined by the sequence of ribonucleotides and environment conditions [5]. It is widely accepted that RNA secondary structure, consisting of canonical Watson-Crick (WC) G-C and A-U and wobble G-U base pairs, forms initially, with other interactions developing subsequently [6]. Since the RNA secondary structure is critical in defining the molecule’s global fold, its knowledge is crucial for determining the RNA 3D structure and function [7]. Despite significant advancements in experimental techniques for RNA structure determination, particularly in chemical probing methods [8] and cryo-electron microscopy (cryo-EM) [9], the structures of many functional RNAs remain unknown [10]. Consequently, the computational problem of RNA secondary structure prediction remains highly relevant.

De novo RNA secondary structure prediction problem involves predicting which pairs of nucleotides will form WC and wobble base pairs. The prediction can be made for a single RNA sequence or a multiple sequence alignment. The most widely used approach for single-sequence prediction is free energy minimization [11], where the structure’s free energy is the sum of its elements’ free energies, and the structure with the minimum free energy (MFE) is identified using dynamic programming [12]. Some methods use probabilistic approaches to predict the structure of maximum expected accuracy (MEA) [13]. For predictions based on sequence alignments, a classic method involves the identification of covarying pairs of residues that correspond to evolutionary conserved base pairs [14]. Several techniques combine covariation analysis with algorithms to find either the MFE [15] or the MEA structure [13, 14, 16]. Recently, deep learning methods have been developed for both single-sequence and alignment-based predictions, demonstrating superior performance compared to traditional tools [17–19].

Despite the diversity of approaches, there is still no definitive solution for RNA secondary structure prediction, particularly for single RNA sequences or sequence alignments comprising too few or too similar RNA sequences for effective covariation analysis [20]. The methods are becoming increasingly complex without notable improvements in accuracy [19]. Most methods ignore pseudoknots [16, 17, 21], which is a significant oversimplification [22]. Additionally, the majority of methods predict only a single structure and are not suitable for RNA molecules that adopt alternative structures [13, 17, 18, 21], with few exceptions [21, 23]. Many tools cannot incorporate structural restraints or chemical probing data in their predictions [13, 14, 16–18]. Only a limited number of tools predict structures formed by multiple RNA sequences [24]. Deep learning methods often suffer from overfitting, lack interpretability, and do not generalize well to RNAs that exhibit previously unknown structures [25, 26].

In this work, we present SQUARNA, a new RNA secondary structure prediction method based on a greedy stem formation model that overcomes the limitations of previous approaches. Benchmarks demonstrate that SQUARNA matches the performance of the best tools for single sequence inputs and significantly outperforms existing methods for sequence alignments.

## RESULTS

### SQUARNA algorithm for the single-sequence prediction

We formulate the de novo RNA secondary structure prediction problem with single-sequence input as a variant of the partial assignment problem [27], wherein two sets of elements serve as input and an optimal matching between certain elements of the two sets constitutes the output. To solve the given problem we devised a greedy stem formation model, see the Methods section for details.

We consider the input ribonucleotide sequence as each of the two sets of elements simultaneously and predict the G-C, A-U, and G-U base pairs as matches between the elements of the two sets (Figure 1A, left). Since the two sets represent the same RNA sequence and RNA does not form hairpins shorter than three residues [28] we restrict our search to base pairs *(i, j)* where *j ≥ i + 4*. Recognizing the interdependence of base pairs [29, 30] and the tendency for base pairs *(i, j)* to co-occur with base pairs *(i + 1, j - 1)* and/or *(i - 1, j + 1)*, we employ scores of stems (consecutive runs of canonical base pairs) as weights for potential matchings to guide the predictions (Figure 1A, middle). The *stem bp-score* is defined as the sum of the scores of its base pairs. For prediction, we employ a greedy two-step approach named SQUARNA (SQUAre RNA). First, the match *(i, j)* of the highest score is selected and marked as a base pair, and then the stem scores are recalculated by removing the incompatible base pairs *(i, k ≠ j)* and *(m ≠ i, j)* from the consideration. This two-step procedure is iteratively repeated until no matches above the defined threshold remain.

**Figure 1.**
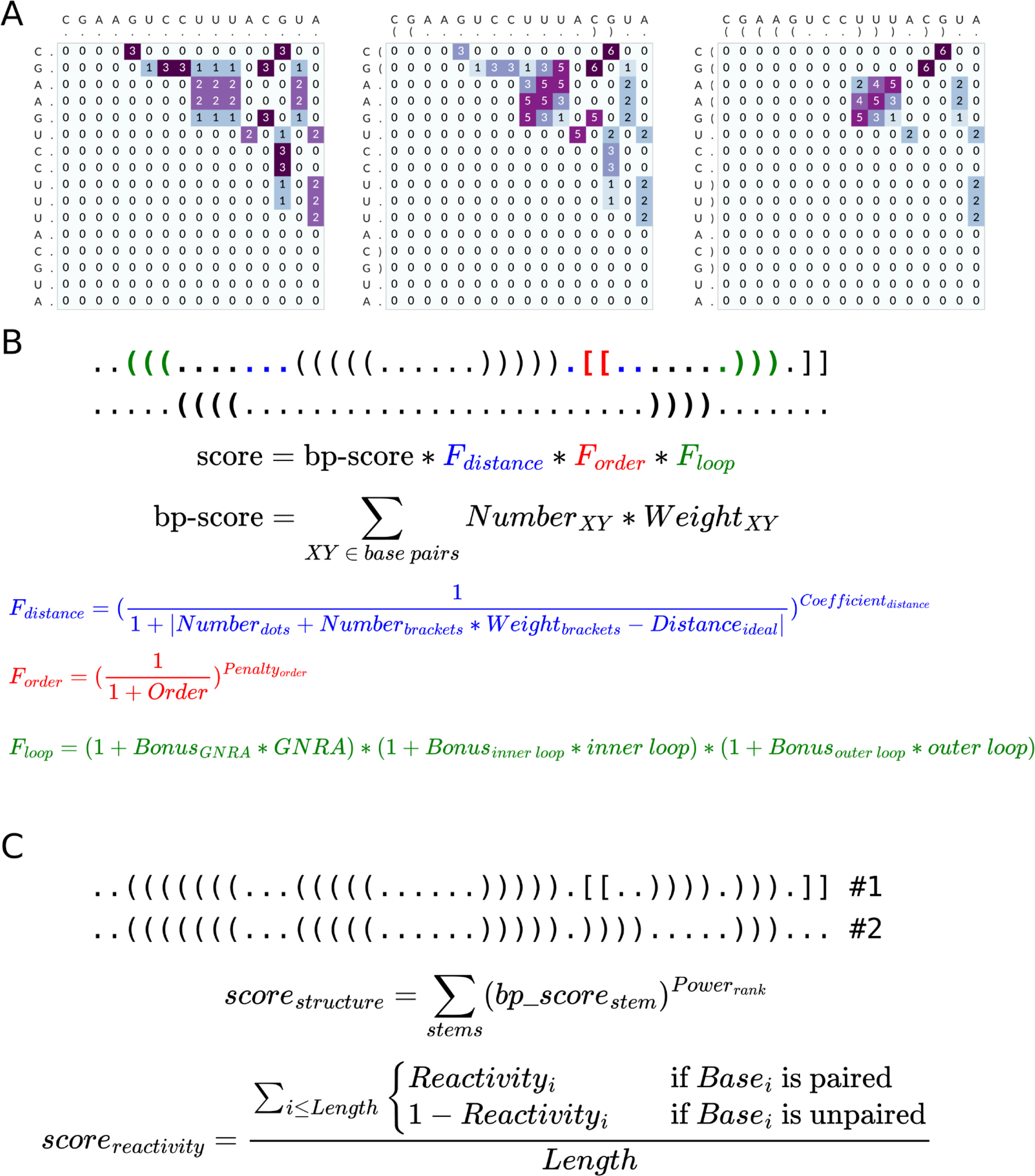
Details of the single-sequence SQUARNA algorithm. (A) An example of solving the partial assignment problem is shown. Left: a square matrix of base pair weights (GC = 3, AU = 2, GU = 1). Middle: the square matrix of stem bp-scores, the stem consisting of the two GC pairs has the highest score; we select the stem and remove the base pairs incompatible with it. Right: the square matrix of stem bp-scores recalculated after the base pair removal, the stem AAG-UUU has the highest score among the remaining stems; we select it and stop the procedure due to having zero other stems of at least two base pairs compatible with the two predicted stems. (B) The stem scoring formula used in SQUARNA. The first dot-bracket line shows the previously predicted stems, and the second dot-bracket line shows the current candidate stem. (C) The structure scoring formulas used for ranking in SQUARNA.

Since all matches of a stem have equal weights, we have reformulated the approach into an equivalent yet more straightforward form with stems being the unit of prediction. In this configuration, we annotate all potential stems, assign scores, select the stem with the highest score, eliminate base pairs incompatible with the chosen stem, and iterate (Figure 1A, middle and right). To better capture the inherent characteristics of RNA secondary structure [28] we use adjusted stem scores (or just *stem scores*) instead of bp-scores. The stem score is calculated as the product of its bp-score and three factors contingent on the previously predicted stems (Figure 1B): a distance factor related to the number of residues confined by the stem, a pseudoknot factor related to the potential pseudoknot’s complexity, and a loop factor providing a bonus for either a GNRA tetraloop or a short near-symmetric internal loop, see the “SQUARNA algorithm for the single-sequence prediction” paragraph in the Methods section for details. If we identify stems that are incompatible with the currently selected stem but have a comparable score (within the defined threshold), we bifurcate and proceed with predictions independently for each such stem added to the current secondary structure. Consequently, SQUARNA generates a set of ranked alternative secondary structures (Figure 1C). SQUARNA can be run with any combination of parameter sets with subsequent merging and overall ranking of the predicted structures.

To benchmark the algorithm’s performance, we compiled the SRtrain dataset of 274 RNA sequences and SRtest dataset of 247 RNA sequences derived from the PDB data bank [31]. The quality of the best out of the top-5 predicted structures was maximized in training procedures, and the *harmonized F-score*, defined as the harmonic mean of the mean F-score among the sequences and the total F-score calculated for the entire number of base pairs in the dataset, was chosen as the target metric. The harmonized F-score was selected to balance the contribution of RNA sequences of different lengths and to reduce the dependence of results on the length distribution in the dataset. Considering the top-5 structures is justified by the RNA 3D structure prediction contests RNA-Puzzles [32] and CASP [33] accepting up to five predictions for each target. For details on the datasets and benchmarks, see the Methods section.

We compared SQUARNA with five state-of-the-art tools (selected based on recent benchmarks [18–20]): three conventional tools, RNAfold/RNAsubopt [21], IPknot [13], and ShapeKnots [23]), and two deep-learning-based tools, MXfold2 [17] and SPOT-RNA [18] (Supplementary Table S1). On the SRtest dataset SQUARNA top-5 performed on the same level as RNAsubopt top-5 and ShapeKnots top-5 with mean F-scores of 82-84% (Supplementary Figure S1, Supplementary Table S2, Supplementary Table S3). Mxfold2, SPOT-RNA, and IPknot produced the best single-structure predictions with mean F-scores of 80-83%. The single-structure predictions of RNAfold, ShapeKnots and SQUARNA demonstrated mean F-scores of 79%, 78%, and 74% respectively.

Therefore, with a much simpler approach, SQUARNA attains the performance of state-of-the-art methods for single-sequence RNA secondary structure prediction.

### SQUARNA algorithm for the alignment-based prediction

For de novo RNA secondary structure prediction from multiple sequence alignment input, we developed a two-step procedure based on the principles of the single-sequence SQUARNA algorithm. Due to unclear stem demarcations in the alignment caused by insertions and deletions, we adopted a base pair as the prediction unit instead of a stem. See the Methods section for details.

Step 1 consists of two iterations (Figure 2). In the first iteration, we calculate matrices of stem bp-scores for each sequence in the alignment (Figure 2A), sum them up to obtain the total stem bp-score matrix (Figure 2B), and then greedily select a subset of compatible base pairs with the highest scores above the defined threshold. In the second iteration, the same procedure is performed but excluding base pairs incompatible with the previously selected subset (Figure 2C). The second iteration serves to refine the predicted structure by eliminating base pairs that belong to mutually incompatible stems with comparable scores. The base pairs selected after the second iteration constitute the predicted structure, referred to as SQUARNA step1.

**Figure 2.**
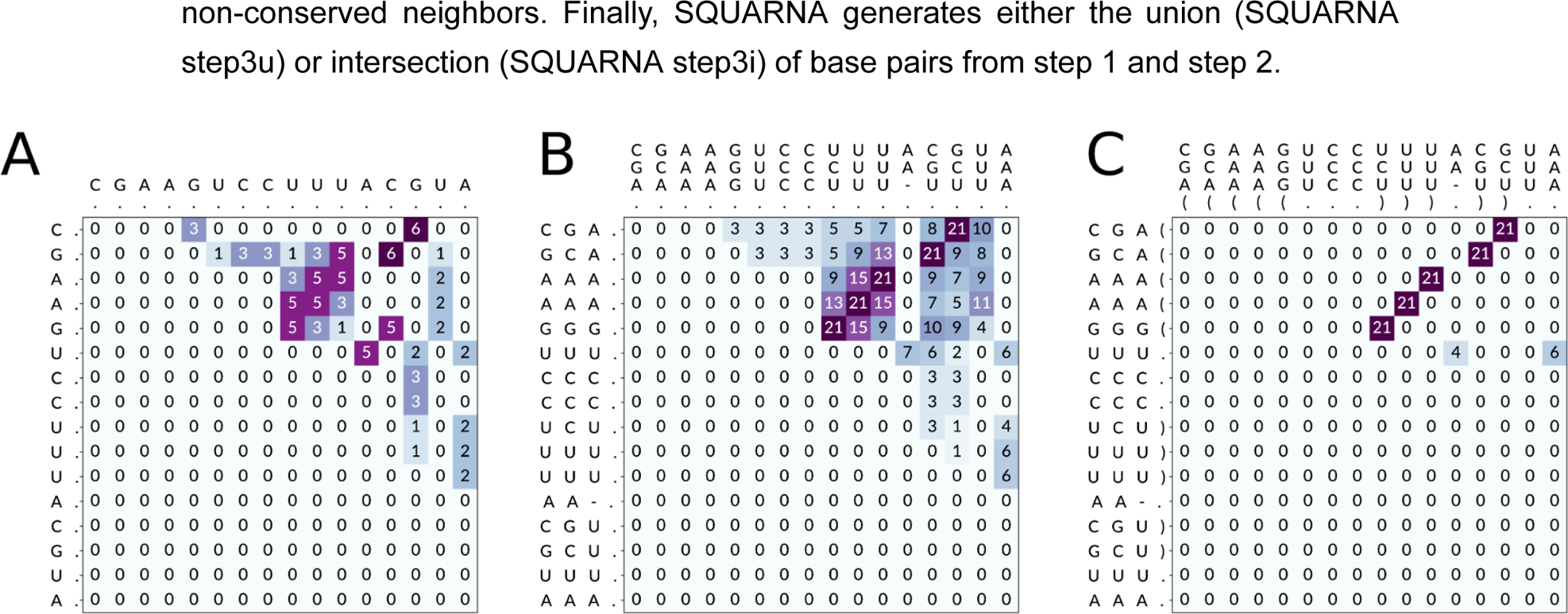
Details of the alignment-based SQUARNA algorithm. Demonstration example of step 1 of the alignment-based SQUARNA algorithm with a score threshold of 10. (A) a square matrix of stem bp-scores (GC = 3, AU = 2, GU = 1) obtained from a single sequence. (B) The total stem bp-score matrix obtained from the sequence alignment at iteration 1; the five cells with a score of 21 compose the restraints for the next iteration. (C) The total stem bp-score matrix obtained at iteration 2; the restraints are shown in dot-bracket notation. The final predicted structure coincides with the restraints in this example.

In step 2, we execute single-sequence SQUARNA predictions guided by the total stem bp-score matrix obtained at step 1, iteration 1, and return the consensus of the individual predictions, referred to as SQUARNA step2. Step 2 helps to identify base pairs that occur in the majority of sequences but don’t reach the score threshold due to non-conserved neighbors. Finally, SQUARNA generates either the union (SQUARNA step3u) or intersection (SQUARNA step3i) of base pairs from step 1 and step 2.

For training, we compiled the RNAStralignExt dataset (Supplementary Table S4) - the set of Rfam seed alignments [10] of eleven non-coding RNA families. This dataset includes eight families from the RNAStralign dataset [34], and three additional families exhibiting well-characterized pseudoknotted structures: HDV ribozyme, SAM riboswitch, and Twister ribozyme. For an independent evaluation, we employed the complete set of Rfam seed alignments (Rfam14.9, 4108 families) and the subset of Rfam families with known 3D structures (RfamPDB, 134 families), excluding the eleven RNAStralignExt families from both sets.

We compared SQUARNA with four state-of-the-art tools (selected based on a recent benchmark [19]): CentroidAlifold [16], RNAalifold [15], IPknot [13], and Rscape (CaCoFold) [14] (Supplementary Table S1, Supplementary Table S5). For Rscape, we evaluated both the nested part of the predicted structure (Rscape nested) and the entire predicted structure (Rscape total). As alignment-based predictions are highly dependent on the alignment depth (number of sequences), we analyzed the results across varying depth thresholds (Supplementary Figure S2). We observed that SQUARNA outperforms the other tools in the range between 100 and 1000 sequences on each of the used datasets. Surprisingly, on RNAStralignExt both CentroidAlifold and RNAalifold showed near zero growth in the quality of their predictions with the increase in depth threshold, while IPknot’s performance even decreased in terms of quality.

To mitigate bias arising from the uneven distribution of alignment depths across Rfam families, we compiled the SubAli dataset of randomly sub-sampled alignments of depths ranging from 2 to 300 sequences for each family from RNAStralignExt. On this dataset, SQUARNA significantly outperformed all other tools (Supplementary Table S6) and only SQUARNA and Rscape demonstrated consistent performance improvement with the increase in depth threshold (Supplementary Figure S2). We further analyzed the results separately for each depth (Figure 3). SQUARNA commenced with a 50% mean F-score on two sequences, reached 80% at ten sequences, and stabilized at 90% on alignments exceeding 100 sequences (Figure 3C, D). CentroidAlifold and RNAalifold started with a mean F-score of 60% on two sequences and plateaued around 75% from five sequences onwards (Figure 3A, D). Rscape exhibited a sharp increase from 30% to an 80-85% mean F-score in the range of 2 to 100 sequences, stabilizing thereafter (Figure 3B, D).

**Figure 3.**
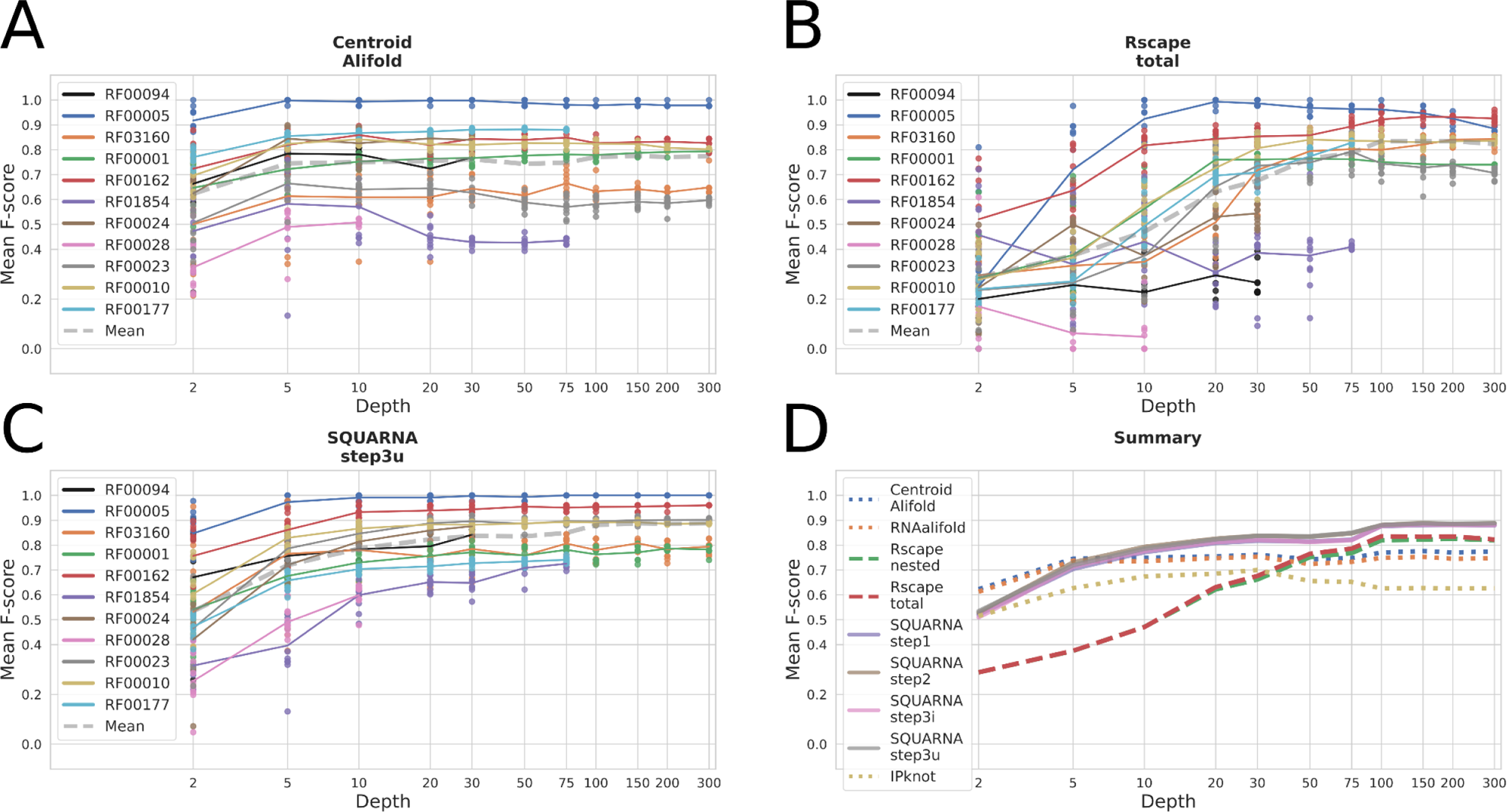
Analysis of the alignment-based prediction quality dependence on alignment depth. (A, B, C) Results of three representative tools (CentroidAlifold, Rscape total, SQUARNA step3u) on the SubAli dataset. Each dot represents the F-score of a single prediction, and solid lines depict the F-scores averaged over the ten alignments of each depth for a specific Rfam family. The dashed gray line indicates the mean F-scores across all eleven Rfam families. (D) Mean F-scores across all eleven Rfam families for each tool against alignment depth.

To assess the influence of sequence similarity on prediction quality, we randomly sampled 5-, 10-, and 20-sequence alignments with varying mean pairwise sequence similarity for the families from RNAStralignExt. Interestingly, only SQUARNA exhibited a subtle yet discernible negative correlation between prediction quality and mean pairwise sequence similarity in the alignment, a correlation that diminished with increasing alignment depth (Figure 4, Supplementary Figure S3).

**Figure 4.**
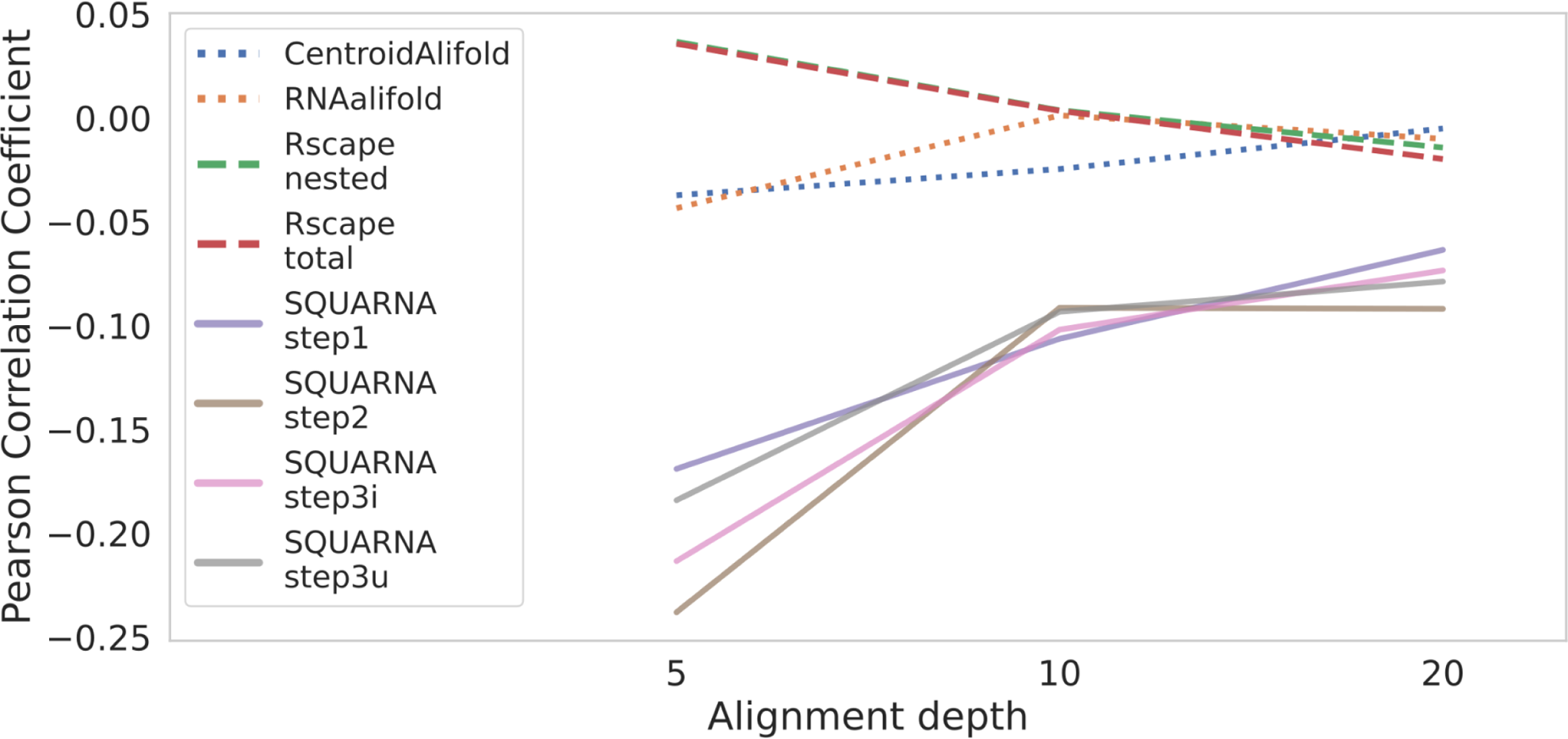
Influence of sequence similarity on prediction quality. The Pearson correlation coefficients between the prediction quality of alignment-based tools and the mean pairwise similarity of the input alignment plotted against the alignment depth. SQUARNA exhibits a noticeable negative correlation, whereas other tools demonstrate values close to zero.

We further examined specific instances where SQUARNA encountered challenges in the deepest alignments of the Rfam14.9 dataset. For the algC RNA motif (RF02929), SQUARNA failed to identify the sole correct stem and instead predicted a two-stem pseudoknot, resulting in a 0.0 F-score despite the alignment depth of 492 sequences (Figure 5A). Upon inspecting the sequences, we discovered that the alignment comprised only 90 unique sequences, with the 3′-half displaying unusually high sequence conservation. This led SQUARNA to identify the pseudoknot, despite the correct stem being the only one to show base pair covariation. However, SQUARNA successfully derived the correct stem from sub-alignments featuring 5 to 40 most diverse sequences of the family. Additionally, the correct stem is discernible in the SQUARNA’s verbose output for the entire seed alignment (Figure 5A).

**Figure 5.**
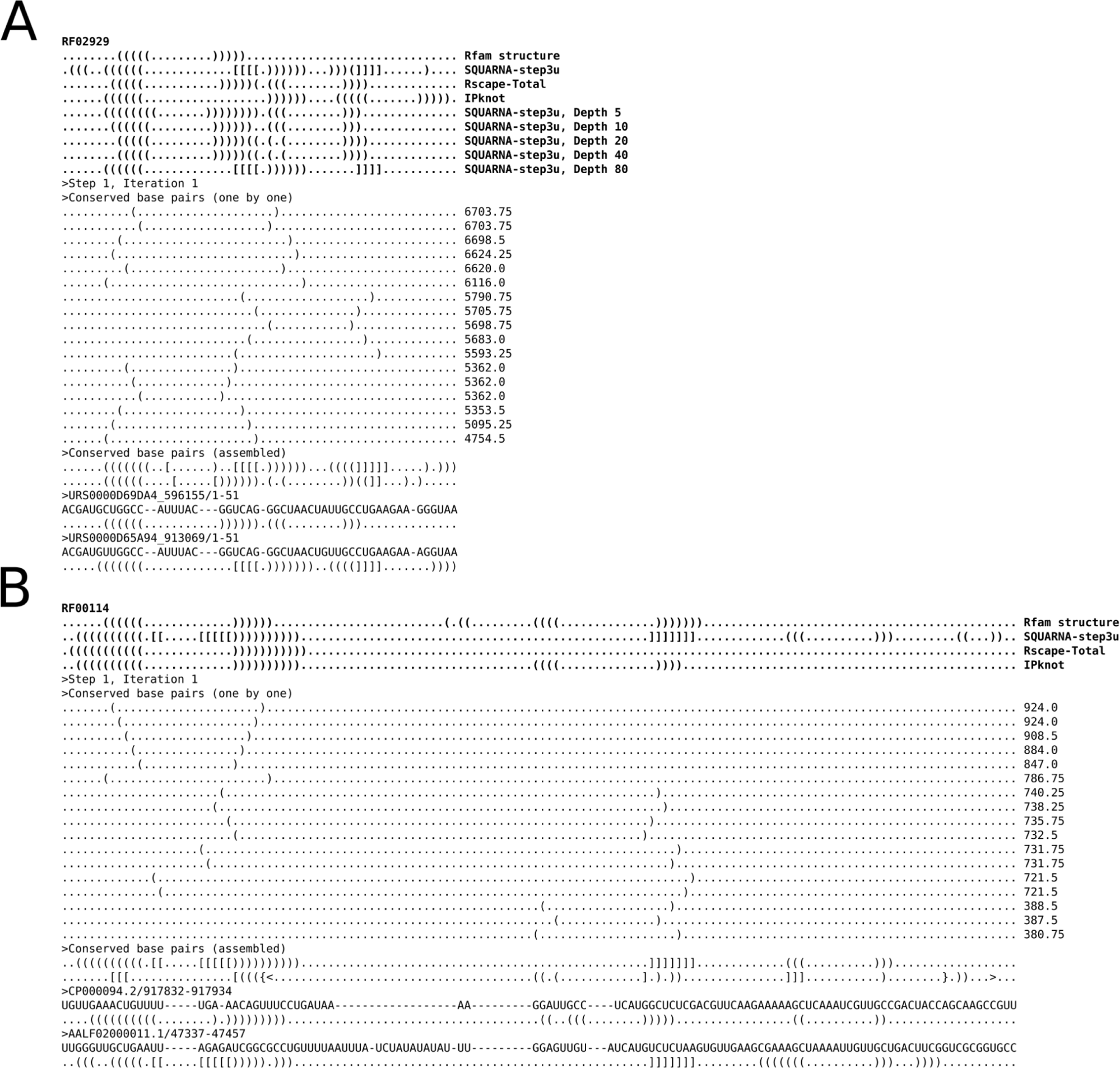
Selected instances showcasing the alignment-based SQUARNA algorithm’s performance. (A) algC RNA motif, Rfam family RF02929, seed alignment of 492 sequences, (B) Ribosomal S15 leader sequence, Rfam family RF00114, seed alignment of 78 sequences. The Rfam consensus structure and different predicted structures are shown in bold at the top of each panel, followed by fragments of the SQUARNA tool verbose output featuring individual sequence predictions at the bottom of each panel.

Another example is the ribosomal S15 leader sequence (RF00114), where SQUARNA achieved a mere 34% F-score on the seed alignment of 78 sequences (Figure 5B). However, the S15 leader sequence is known to adopt two alternative folds: one involving consecutive hairpins and another featuring a pseudoknot [35]. The Rfam structure includes the two hairpins, while SQUARNA was the only tool to correctly identify the pseudoknot. Moreover, the second hairpin can be observed in the SQUARNA’s verbose output (Figure 5B).

As the S15 leader sequence example shows, the secondary structures currently annotated in Rfam are not perfect to serve as ground truth. Therefore, as an additional test, we also assessed the number of base pairs predicted by SQUARNA for the Rfam seed alignments of the Rfam14.9 dataset that were also identified as significantly covarying by Rscape [14]. SQUARNA’s predictions included 5633 covarying base pairs, which is just 13 base pairs less than those found to co-vary by Rscape in the Rfam-annotated secondary structures (Supplementary Table S7). Secondary structures predicted by SQUARNA are well-supported by Rscape in terms of significantly covarying base pairs as the Rfam annotations, better than predictions of other methods (Supplementary Figure S4).

### Chemical probing input data

To assess the impact of chemical probing data on prediction quality from the final user perspective, we prepared the S01 dataset of 24 challenging RNAs [23] in four different settings: (i) RNA sequence only (sequence); (ii) RNA sequence and SHAPE data (sequence+shape); (iii) alignment of homologous sequences (alignment); (iv) alignment of homologous sequences and SHAPE data (alignment+shape). The multiple sequence alignments were prepared using a structure-unaware approach, involving BlastN [36] for homolog search and MAFFT [37] for alignment of the obtained hits.

SQUARNA with the “sk” parameter set (SQUARNAsk top-5) exhibited a mean F-score of 71% with sequence input, improving to 82% with the addition of SHAPE data (second-best result after 88.5% exhibited by ShapeKnots, see Supplementary Figure S5 and Supplementary Table S8). Notably, SQUARNA performed particularly well on shorter sequences, achieving the highest mean F-score of 93% for the 14 sequences under 200 nucleotides using sequence+shape data, while ShapeKnots-top-5 achieved 91% for this subset. For alignment+shape input, SQUARNA demonstrated a 2.5-4.5% improvement compared to alignment input. However, overall, the prediction quality of the alignment-based tools was considerably lower than that of the single-sequence tools. Surprisingly, predictions on structural alignments built by covariation models [38] still exhibited lower accuracy than single-sequence predictions (Supplementary Figure S6, Supplementary Table S9).

On the Ribonanza dataset, which was not used for training SQUARNA or any other existing tools, SQUARNA emerged as the sole tool capable of effectively utilizing DMS [39] and 2A3 [40] input reactivity data, resulting in an impressive mean F-score of 85% (Figure 6, Supplementary Table S10). For the details on the Ribonanza dataset, refer to the Methods section.

**Figure 6.**
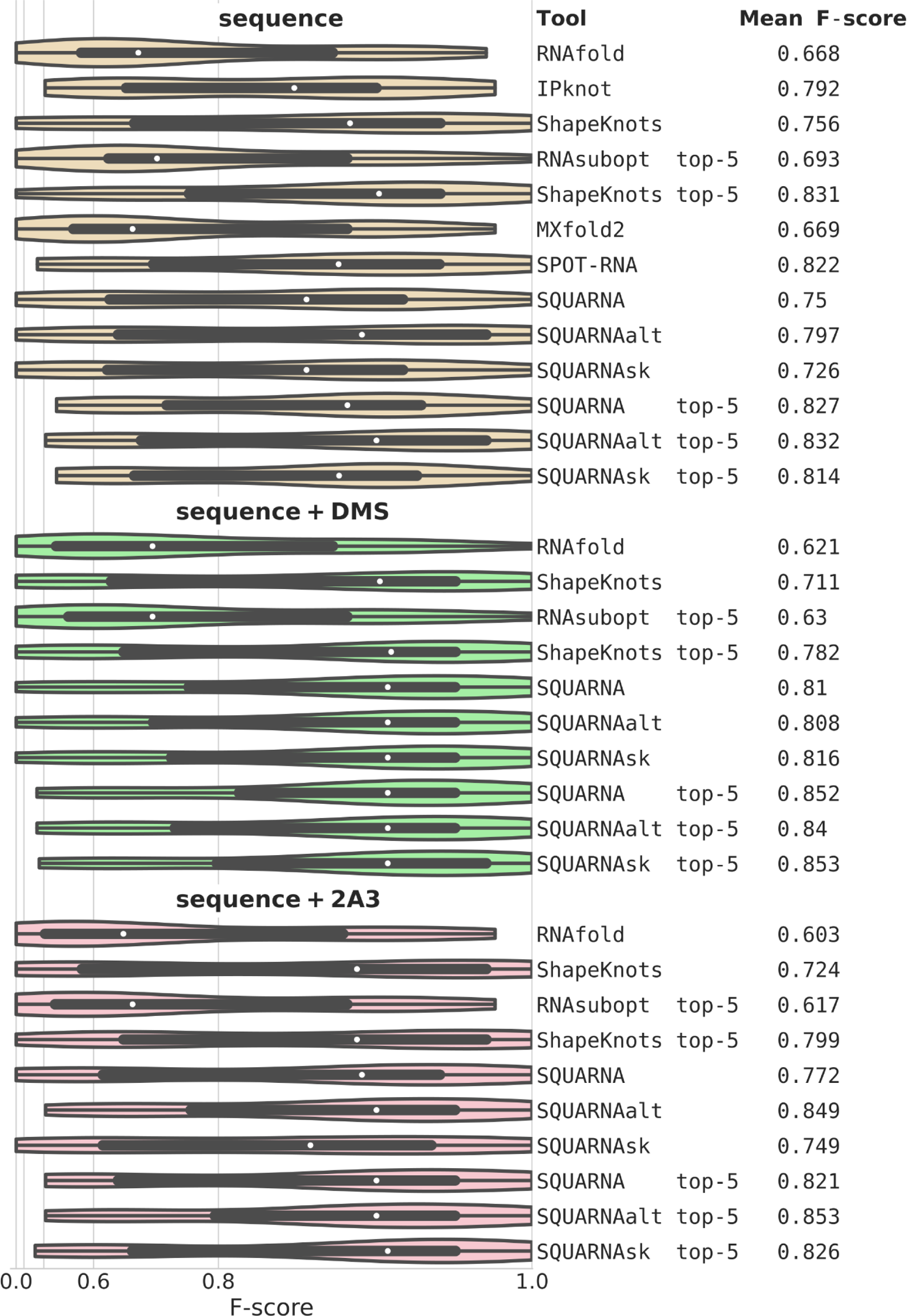
Benchmarking results with chemical probing data. Benchmark results on the Ribonanza dataset comprising 22 sequences ranging from 29 to 95 nucleotides in length. The violin plots depict in exponential scale F-score distributions derived from predictions based on three different input types: sequence, sequence with DMS reactivities data, and sequence with 2A3 reactivities data. Boxplots are provided within the violins, with white dots denoting median values. Mean F-scores are listed on the right side of the figure. The graph was generated using the Seaborn Python library (https://seaborn.pydata.org/generated/seaborn.violinplot.html).

## DISCUSSION

In this work, we introduced SQUARNA, a new approach to de novo RNA secondary structure prediction employing a stem formation model. Our benchmarks show that the single-sequence SQUARNA performs on par with state-of-the-art methods, while the alignment-based SQUARNA significantly outperforms the existing tools.

The greedy approach employed in SQUARNA is meant to mimic RNA folding kinetics and, crucially, circumvents global optimization procedures impeded by pseudoknots, such as free energy minimization. Conveniently, the first-predicted stems are the most confidently predicted (Supplementary Figure S7), enabling fine-tuning of the precision/recall ratio by restricting the number of stems to be predicted by SQUARNA. Additionally, the adoption of stem scores over base pair scores enhances the “signal to noise ratio”, allowing alignment-based SQUARNA to attain the CentroidAlifold/RNAalifold baseline on as few as five sequences, while Rscape requires around 50 sequences to achieve a comparable result (Figure 3D). Notably, a significant portion (46.5%) of Rfam seed alignments comprise 5 to 50 sequences, whereas only 7.7% of the alignments feature over 50 sequences, sufficient for Rscape to confidently predict a structure. Furthermore, SQUARNA step1 ranks as the second-fastest alignment-based tool after RNAalifold. Nevertheless, SQUARNA’s key limitation is its susceptibility to being influenced by sequence conservation over covariation, a challenge mitigated through careful curation of the input alignment.

Single-sequence SQUARNA reached the level of the leading deep-learning methods while employing a much simpler approach devoid of the typical flaws inherent to learning-based algorithms, such as overfitting, lack of interpretability, and poor generalization to unfamiliar data [25]. SQUARNA effectively handles any natural or synthetic RNA sequence, and it even allows users to track the sequential prediction of stems. Moreover, SQUARNA seamlessly processes any type of modified RNA bases, whether common or rare [41], provided the user specifies base pair weights for such bases. The harmonized F-score metric employed in this study for benchmarking revealed that certain tools, particularly SPOT-RNA and SQUARNA, perform better for shorter RNA sequences (Supplementary Table S2). This can be addressed in the future by developing distinct SQUARNA settings tailored to different RNA sequence length ranges.

SQUARNA can successfully utilize both RNA single sequence and sequence alignment inputs as well as chemical probing data (Figure 6). Additionally, SQUARNA can detect alternating folds (Figure 5). Moreover, SQUARNA can incorporate structural restraints as supplementary input and predict RNA secondary structure formed by multiple RNA sequences, allowing users to assess the oligomerization potential of a given RNA. We are confident that the inherent strengths of SQUARNA will profoundly impact the field of RNA computational biology.

## METHODS

### Theoretical background

The SQUARNA approach is rooted in the ARTEM algorithm, designed for the superposition of two arbitrary RNA 3D structure fragments without prior knowledge of nucleotide matchings between the fragments [42]. The two core ideas of ARTEM are the formulation of the problem as the partial assignment problem and the utilization of dependencies in the data. In the context of de novo RNA secondary structure prediction, the partial assignment pertains to the base pairs formed between nucleotides, and the primary data dependency is the tendency of base pairs to form continuous stems.

In the initial phase of our preliminary analysis, we observed optimal results with negative weights for GU base pairs. This introduced an ambiguity in defining a stem: as either the longest sequence of consecutive base pairs or as the subsequence of consecutive base pairs with the highest bp-score. The stems from the known RNA secondary structures were found to be predominantly the stems of maximum length, with only a few instances where one flanking GU/AU base pair was absent from the reference structure stems (data not shown). We tested all possible stem definitions, including the definition of a stem as any possible subsequence of consecutive base pairs. However, the results remained largely unchanged. Consequently, we ultimately defined a stem as the longest possible sequence of consecutive base pairs, as it proved to be the most efficient for the algorithm runtime.

For the description of RNA secondary structure elements, such as loops and pseudoknots, we used the definitions from [43]. *Pseudoknot order* (or *pseudoknot complexity*) was defined as the minimum number of different bracket types required for its non-ambiguous representation in dot-bracket format. Bracket type assignment for the base pairs of a given structure was performed greedily, followed by a bubble-like sorting to minimize the number of higher-order base pairs, see the PairsToDBN function in the code: https://github.com/febos/SQUARNA/blob/main/SQRNdbnseq.py. Two base pairs *(i, j)* and *(k, l)* were considered *inconsistent* (*intersecting*) with each other if they couldn’t be formed simultaneously (i.e., if *i = k* or *i = l* or *j = k* or *j = l*), and *consistent* (*non-intersecting*) otherwise. Two stems were considered *consistent* with each other if all their base pairs were consistent, otherwise, the stems were considered *intersecting*.

### RNA sequence datasets

The SRtrain dataset was prepared as follows. RNA 3D structures from the PDB [31] were selected according to a representative set of RNA structures [44] (version 3.278 with a 3.0 Å cutoff). RNA secondary structures were annotated using DSSR [45] (version 2.0). Subsequently, a custom Python script was used to extract RNA strands from the PDB entries, meeting three conditions: (i) the strand is continuous, devoid of chain breaks or missing residues; (ii) the strand does not involve modified residues; (iii) the strand does not form WC or wobble base pairs with any other strand but forms at least one such base pair within the strand. The set of derived strands underwent manual secondary structure redundancy reduction, resulting in a non-redundant subset of 274 RNA strands with their secondary structures. The SRtrain150 subset included 266 strands with a length shorter than 150 nts.

The SRtest dataset was collected using the same procedure from another version of the representative set of RNA structures [44] (version 3.322 with no resolution cutoff). Sequences with at least 80% identity to any sequence from the SRtrain dataset were excluded. The resulting set of derived strands underwent manual secondary structure redundancy reduction, yielding a non-redundant subset of 247 RNA strands with their secondary structures.

The TS1reducedWC dataset was prepared as follows. The set of 62 RNA sequences populating the TS1reduced dataset was derived from [18] and converted from .ct format to dot-bracket format. However, the dataset contained a mix of canonical and non-canonical base pairs without specific annotation. Consequently, the original secondary structures were not used; instead, structures for the 62 sequences were derived using DSSR (version 2.0) to ensure inclusion of only WC and wobble base pairs. The resulting dataset was named TS1reducedWC.

The S01 dataset was prepared as follows. The set of 24 sequences along with SHAPE data, was downloaded from [23] (https://webshare.oasis.unc.edu/weeksgroup/data-files/ShapeKnots_DATA.zip). RNA secondary structure data were converted from .ct format to dot-bracket. The original SHAPE data contained values ranging from 0.0 to 1.5, with some outliers and missing values set to −999. Two sequences required adjustments: (i) for the entry “5’ domain of 16S rRNA, H. volcanii” one excess value from the 3’-end was removed; (ii) for the entry “SARS corona virus pseudoknot” three missing −999 values were added at the 3’-end. The supplementary information from [23] (https://www.pnas.org/doi/suppl/10.1073/pnas.1219988110/suppl_file/sd01.pdf) served as a reference for these fixes. For predictions with ShapeKnots and ShapeSorter, the original SHAPE data (after the fixes) were used. For predictions with RNAfold, RNAsubopt, and RNAalifold, the values were scaled to range from 0.0 to 2.0 (divided by 1.5 and multiplied by 2.0), outliers were truncated (values under 0.0 were set to 0.0, and values higher than 2.0 were set to 2.0), and missing values were kept as −999. For predictions with SQUARNA, the values were similarly scaled to range from 0.0 to 1.0, outliers were truncated, and missing values were set to 0.5.

The Ribonanza dataset was prepared as follows. The DMS and 2A3 reactivity data were derived from the training set of the Ribonanza competition at Kaggle [46]. Raw reactivity values were clipped to the range from 0.0 to 1.0 (i.e., outliers were truncated) and missing values along with DMS values for G and U were set to −999. Next, a BlastN search for RNA chains of at least 20 nucleotides from the PDB against the competition training set was conducted. Identical hits with 100% query coverage were selected. Among the RNA chains with at least one hit, continuous strands forming isolated secondary structures, akin to the approach for the SRtrain dataset, were further selected. For the chosen strands, reactivity values were averaged among hits with “SNfilter” = 1, separately for the “DMS_MaP” and “2A3_MaP” experiment types. The resulted dataset comprised 22 RNA sequences from 29 nts to 95 nts in length, with 13 out of the 22 RNAs forming pseudoknots. For predictions with SQUARNA, −999 values were set to 0.5. For predictions with ShapeKnots, values were scaled to range from 0.0 to 1.5. For predictions with RNAfold and RNAsubopt, values were scaled to range from 0.0 to 2.0. Homologous sequence alignments were not prepared for the Ribonanza dataset, as BlastN couldn’t find at least two homologs for 17 of the 22 RNA sequences.

For each of the prepared datasets, NL (No-Lone) versions were generated (SRtrainNL, TS1reducedWCNL, etc.), with the lone base pairs removed. However, preliminary experiments indicated minimal differences in benchmarks between the standard and NL versions of the datasets. Consequently, the NL benchmarks were eventually omitted.

### RNA multiple sequence alignment datasets

The RNAStralignExt, RfamPDB, and Rfam14.9 datasets were prepared by obtaining seed alignments in Stockholm format (https://en.wikipedia.org/wiki/Stockholm_format) from Rfam [10] (version 14.9, 4108 families). The Stockholm files were then converted into one-line Fasta-like (https://en.wikipedia.org/wiki/FASTA_format) and Clustal (https://meme-suite.org/meme/doc/clustalw-format.html) formats with custom Python scripts. The 134 families of the RfamPDB dataset were identified via a search for “Families with 3D structure” on the Rfam web page (https://rfam.org/search?q=entry_type:%22Family%22%20AND%20has_3d_structure:%22Yes%22). All benchmarks for Rfam14.9 and RfamPDB were conducted with the exclusion of the eleven RNAStralignExt families.

The SubAli dataset was prepared as follows. For each of the eleven Rfam families of the RNAStralignExt dataset, ten alignments were randomly sampled at each of the eleven depths (where the seed alignment depth allowed it): 2, 5, 10, 20, 30, 50, 75, 100, 150, 200, and 300 sequences. The sampled sequences were not subjected to any realigning procedures.

The SeqSim dataset, used to analyze the correlation between prediction quality and mean sequence similarity of the alignment, was prepared as follows. For each of the eleven Rfam families of the RNAStralignExt dataset, 100 alignments were randomly sampled at each of the three depths - 5, 10, and 20 sequences (where the seed alignment depth allowed it). For each alignment of a given (family, depth) pair, the mean pairwise sequence similarity was calculated and rounded to three decimal digits. Subsequently, only one alignment for each (family, depth, similarity) triple was retained, resulting in approximately 50 different alignments for a (family, depth) pair. The Hamming distance (https://en.wikipedia.org/wiki/Hamming_distance) between the aligned sequences was used as the measure of pairwise sequence similarity. The sampled sequences were not subjected to any realigning procedures.

The S01AliUngap dataset, used to evaluate prediction quality on the S01 dataset with structure-unaware alignment and alignment+shape input, was prepared as follows. For each of the 24 sequences, BlastN [36] (https://blast.ncbi.nlm.nih.gov/Blast.cgi?PROGRAM=blastn&PAGE_TYPE=BlastSearch&LINK_LOC=blasthome) was used to search for homologs with at least 90% query sequence coverage. Unique homologs were derived from the search results. If the resulting number of sequences exceeded 100, the 100 most diverse sequences were selected as follows. First, the reference sequence was chosen. Then, sequences were iteratively added one by one until reaching 100 sequences. At each step, the sequence with the lowest highest similarity to the previously selected sequences was chosen. The resulting set of sequences was realigned with MAFFT [37] (version v7.453). Columns with gaps in the reference sequence were then removed.

The S01AliCM dataset, used to measure prediction quality on the S01 dataset with structure-aware alignment and alignment+shape input, was prepared similarly to S01AliUngap, except with the use of Infernal [38] (v.1.1.4) instead of MAFFT. Covariation models were prepared based on the reference sequences along with the nested parts of the reference secondary structures using *cmbuild.* The homologous sequences were then aligned using *cmalign*. Columns with gaps in the reference sequence were subsequently removed.

The sub-alignments of the most diverse sequences of the RF02929 family were prepared as follows. First, the two most dissimilar sequences were selected. Subsequently, new sequences were iteratively added one by one until reaching the required depth. At each step, the sequence with the lowest highest similarity to the previously selected sequences was chosen.

### Quality metrics

To assess prediction quality, the number of true positives (*TP*) was defined as the number of correctly predicted base pairs, the number of false positives (*FP*) as the number of wrongly predicted base pairs, and the number of false negatives (*FN*) as the number of wrongly missed base pairs. We avoided defining the number of true negatives (*TN*). The precision metric was defined as *Precision = TP / (TP + FP)*, the recall metric was defined as *Recall = TP / (TP + FN)*, and the F-score metric was defined as the harmonic mean of precision and recall, *F-score = 2 * Precision * Recall / (Precision + Recall) = 2 * TP / (2 * TP + FP + FN)*. In addition to the commonly used mean F-score value among the sequences of a given dataset, we used the total F-score value, calculated from the *TP*, *FP*, and *FN* values summed up over the sequences of a given dataset. Finally, the harmonized F-score was defined as the harmonic mean between the mean F-score and total F-score values, serving as the main target metric.

The statistical significance of the differences in the tools’ predictions was calculated using Student’s paired t-test [47] as implemented in *ttest_rel* function from scipy.stats python3 library

(https://docs.scipy.org/doc/scipy/reference/generated/scipy.stats.ttest_rel.html).

### Other tools used for benchmarking

The RNAfold, RNAsubopt, and RNAalifold tools were locally installed as part of the ViennaRNA package (version 2.5.0) and executed with the following commands: “RNAfold --noPS [--shape=shapefile] < inputfile > outputfile”, “RNAsubopt --sorted [--shape=shapefile] < inputfile > outputfile”, and “RNAalifold --noPS inputfile [--shape=shapefile] > outputfile”. Due to RAM limitations, the “--deltaEnergy=0.1” setting was used to run RNAsubopt for sequences longer than 2000 nts. IPknot, obtained from https://github.com/satoken/ipknot (version 1.0.0), was installed locally and executed with the command: “ipknot inputfile > outputfile”. However, on the SeqSim dataset, IPknot exceeded ten days of execution on the alignment RF00177_5_729.sto, and on the Rfam14.9 dataset, it exceeded ten days on the RF02746 family alignment. Consequently, IPknot benchmarks for these two datasets could not be obtained. ShapeKnots, part of the RNAstructure package (version 6.4), was installed locally and run using the command: “DATAPATH=../data_tables./ShapeKnots-smp inputfile outputfile [-sh shapefile]”. MXfold2, obtained from https://github.com/mxfold/mxfold2 (version 0.1.2), was installed locally and executed with the command: “mxfold2 predict inputfile > outputfile”. SPOT-RNA, from https://github.com/jaswindersingh2/SPOT-RNA (as of April 16, 2023) was installed locally and run using the command: “python SPOT-RNA.py --inputs inputfile --outputs outputfolder --cpu 32”.

CentroidAlifold, obtained from https://github.com/satoken/centroid-rna-package (version v0.0.17), was installed locally and run with the command: “centroid_alifold inputfile > outputfile”. Rscape, available at http://eddylab.org/R-scape/ (version 1.6.1), was installed locally and run for prediction with the command: “R-scape --fold --covmin 4 --nofigures --rna inputfile > outputfile”. The “Rscape nested” structure was defined from the “SS_cons” line of the output, and the “Rscape total” structure was defined from the “SS_cons” line with the addition of consistent base pairs from the “SS_cons_N” lines of the output. To identify significantly covarying base pairs we run Rscape with the command “R-scape --nofigures --rna inputfile > outputfile” and used the default E-value threshold of 0.05. ShapeSorter predictions were obtained using the web server version at https://e-rna.org/shapesorter/ (as of August 22, 2023), as we couldn’t run it locally. As ShapeSorter predicts separate stems, the predicted structures were built by greedily selecting the predicted stems below the recommended p-value threshold of *7 * 10^−4^* [48]. SPOT-RNA2 [19] was not included in the benchmarks as it takes a single-sequence input and builds an alignment of homologous sequences on the fly. Consequently, it’s unclear to which category of the tools it should belong.

The benchmarks were conducted on an AMD Ryzen 9 5950X machine equipped with 32 CPU cores and 128 GB RAM.

### SQUARNA algorithm for the single-sequence prediction

The single-sequence SQUARNA algorithm comprises the following steps:

0) Initialization with an empty secondary structure as the only currently predicted structure;
1) annotation of all possible stems above the defined minimum length (minlen) and minimum bp-score (minbpscore) thresholds that are consistent with (but not a part of) the currently predicted structure;
2) calculation of stem scores according to the stem score formula (Figure 1B);
3) removal of the stems with scores below the threshold defined as the product of the minbpscore and a scaling factor (minfinscorefactor);
4) sorting the stems in decreasing order based on their stem scores;
5) selection of the stem with the highest score along with all the intersecting stems with a score within the suboptimality threshold (subopt);
6) adding the selected *k* stems to the currently predicted structure separately, one by one, resulting in *k* new currently predicted structures;
7) repeat steps 1-6 for each of the currently predicted structures in the current pool and obtain a new pool. Repeat until the pool of currently predicted structures is empty. If, for a given currently predicted structure, the maximum allowed number of stems is reached (maxstemnum) or no stem can be added to the structure, the structure is moved from the pool to the set of predicted structures;
8) repeat steps 0-7 for each of the defined parameter sets;
9) all the predicted structures are ranked by their structure score, reactivity score, or the product of the two scores (Figure 1C).

The *subopt* parameter is initialized with the starting value (suboptmin) and is increased by an increment value each time the size of the pool of currently predicted structures grows (on step 7) until the maximum defined value is reached (suboptmax). The increment value is determined by the number of increment steps (suboptsteps) as *(supoptmax - suboptmin) / suboptsteps*.

Any hard restraints defined by the user are considered in step 1, allowing only the base pairs consistent with the restraints. In the case of multi-chain input, the distance factor is always set to 1 for the stems confining the chain breaks, and *(i, i + 1)*, *(i, i + 2)*, and *(i, i + 3)* base pairs are allowed for such stems. In the case of reactivities defined by the user, *(i, j)* base pairs of positive weights are multiplied by, and negative weights are divided by a reactivity factor defined as *(1 - (reactivity_i_ + reactivity_j_) / 2) * 2*. The reactivities are expected to range from 0.0 to 1.0, with 0.5 being the neutral value, but outliers will be handled as well.

The theoretical time complexity of the algorithm is *O(log(K) * N^4^)*, where N is the sequence length and K is the number of predicted structures. Step 1 takes *O(N^2^)* time to annotate all the stems, step 2 takes *O(N)* time to score each of the *O(N^2^)* annotated stems, resulting in *O(N^3^)* time. The number of times steps 1-6 are repeated is equal to the *O(N)* number of stems in a predicted structure, resulting in *O(N^4^)*. Overall, the algorithm runs relatively fast with *subopt = 1.0*, but significantly slows down with lower *subopt* values as the number of alternatives *K* grows exponentially.

The initial training procedure was performed on the SRtrain150 dataset in the form of a sequence of grid-search runs. The target metric was the harmonized F-score of the best of the top-5 structures. After training, the following fixed values were chosen: *Distance_ideal_* was set to 4 for hairpins and 2 for all other loops (distance factor formula in Figure 1B); the internal loops of the following sizes were defined as short near-symmetric loops deserving a bonus: *(0, 0), (0, 1), (1, 0), (1, 1), (0, 2), (2, 0), (2, 2), (1, 2), (2, 1), (3, 1), (1, 3), (2, 3), (3, 2), (3, 3), (3, 4), (4, 3), (4, 4), (4, 2), (2, 4)*; for symmetric loops *(x, x)* the bonus was doubled; for *(x - 1, x)* and *(x, x - 1)* loops the bonus was multiplied by 1.5; *Bonus_GNRA_* was set to 1.25; for the structure score (Figure 1C) the bp weights were set to *GU = −1.0, AU = 1.5, GC = 4.0*, and the *Power_rank_* was set to 1.7. No bifurcations were allowed after the size of the pool of currently predicted structures reached 1000 structures.

After training, the following parameter values were chosen: the ranking is performed based on the structure score (this was set to default and used for all the benchmarks); the default config (*def.conf*) included two parameter sets - *def1* and *def2*; the *def1* parameter set included the following values: bp weights *GC = 3.25, AU = 1.25, GU = −1.25*; *suboptmax = 0.9*; *suboptmin = 0.65; suboptsteps = 1; minlen = 2; minbpscore = 4.5; minfinscorefactor = 1.25; Coefficient_distance_ (distcoef) = 0.09; Weight_brackets_ (bracketweight) = −2; Penalty_order_ (orderpenalty) = 1.0; Bonus_inner loop_ & Bonus _outer loop_ (loopbonus) = 0.125;* the maximum number of stems to be predicted was not limited (*maxstemnum = 1e6*); the *def2* parameter set differed from *def1* in the following values: bp weights *GC = 2, AU = 1, GU = 1; minbpscore = 3; minfinscorefactor = 0.99; distcoef = 0.1; orderpenalty = 1.35*. An alternative config (*alt.conf*) with the best single parameter set “*alt*” was saved, which also works relatively better for short pseudoknotted RNAs. The *alt* parameter set differed from *def1* in a single value: *minfinscorefactor = 1.0*.

The *sk* config (*sk.conf*) was trained on a subset of the S01 dataset involving 14 sequences shorter than 200 nts along with their SHAPE data and included two parameter sets - *sk1* and *sk2*. The *sk1* parameter set differed from the *def1* set in the following values: *suboptmin = 0.75; suboptsteps = 2; minbpscore = 7; orderpenalty = 0.75*. The *sk2* parameter set differed from the *sk1* set in the following values: *bp weights GC = 2, AU = 1, GU = 1; minbpscore = 3; minfinscorefactor = 0.99; distcoef = 0.1; orderpenalty = 1.35*.

Due to the slow performance of the trained SQUARNA configs on longer RNA sequences, two faster configs were designed, one for sequences longer than 500 nts (*500.conf*) and another for sequences longer than 1000 nts (*1000.conf*). The configs were used for sequences of appropriate lengths in all the single-sequence benchmarks. The 500 config included two parameter sets *long1* and *long2* that differed from the *def1* and *def2* sets only in the two values: *suboptmax = 0.95; suboptmin = 0.9*. The 1000 config included a single parameter set that differed from the *def1* set in the two values as well: *suboptmax = 0.99; suboptmin = 0.99*.

### SQUARNA algorithm for the alignment-based prediction

The alignment-based SQUARNA algorithm works as described in the Results section. To calculate the total stem bp-score matrix on step 1, each individual sequence is un-aligned by removing all the gaps. An individual stem bp-score matrix is then calculated based on three parameters (*bpweights, minlen, minbpscore*). After that, the individual matrix is re-aligned to the alignment size by introducing all-zero rows and columns in gap positions of the aligned sequence. An individual structure of a given sequence at step 2 is defined as the consensus of the top-N (1 by default) ranked predicted structures based on a defined config. The consensus structure at step 2 is calculated by greedy selection of a consistent set of the most populated (i.e., present in the highest number of sequences) base pairs from the individual structures present in at least *freqlim* share of the sequences. For each of the step 1 and step 2 predicted structures separately, a trimming procedure is applied, removing the base pairs of pseudoknot orders higher than the defined threshold (*levellim*). On step 3, the union of the step 1 and step 2 base pairs (in the *step3u* setting) is calculated by adding consistent step 2 base pairs to the base pairs obtained after step 1.

The parameter training procedure was performed on the RNAStralignExt dataset. The target metric was the mean F-score of the SQUARNA *step3u* setting. After the training, the following parameter values were chosen: *levellim = 3* for alignment length under 500 nts and 2 otherwise; *freqlim* was set to 0.35; step1 parameters: *bp weights GC = 3.25, AU = 1.25, GU = −1.25, minlen = 2, minbpscore = 4.5*; the config for the single-sequence SQUARNA predictions at step 2 (*ali.conf*) involved a single parameter set “*ali*” with the following values: *bp weights GC = 3.25, AU = 1.25, GU = −1.25; suboptmax = 1.0; suboptmin = 1.0; suboptsteps = 1; minlen = 2; minbpscore = 4.5; minfinscorefactor = 1.0; distcoef = 0.09; bracketweight = −2; orderpenalty = 0.75; loopbonus = 0.125; maxstemnum = 1e6*. At step 2, base pair scores in the individual predictions were weighted with the total stem bp-score matrix obtained at iteration 1 of step 1. The matrix was normalized to have the maximum value of 1.0 (via dividing all the values by the maximum value). Then, all the matrix values were multiplied by 5.0. The resulting matrix was un-aligned for each individual sequence separately and used to weight the base pair scores.

The theoretical time complexity of SQUARNA step 1 is *O(D * N^2^)*, where *D* is the alignment depth and *N* is the alignment length, as the calculation of *D* individual stem bp-score matrices takes O(N^2^) time to annotate all potential stems. In the benchmarks, SQUARNA step1 was the second fastest algorithm after RNAalifold. The theoretical time complexity of SQUARNA step 2 is *O(D * log(K) * N^4^)*, which is the complexity of the single-sequence SQUARNA multiplied by the number of sequences in the alignment.

### SQUARNA tool implementation

The command-line SQUARNA tool, implemented in Python3, utilizes the multiprocessing module from the Python standard library and is available at https://github.com/febos/SQUARNA. For the single-sequence SQUARNA, parallelization is achieved through the pool of the currently predicted structures. In contrast, alignment-based SQUARNA is parallelized by individual sequences. The tool accommodates various input file formats, including FASTA, Stockholm, and Clustal, along with its own fasta-like format, enabling the incorporation of restraints, reactivities, and reference structure input data. When conducting single-sequence predictions with alignment input, the predicted structures are aligned to the input sequences.

## DATA AVAILABILITY

The benchmarking data are available at https://github.com/febos/SQUARNA (DOI: 10.5281/zenodo.8292325).

## CODE AVAILABILITY

An implementation of SQUARNA tool is available at https://github.com/febos/SQUARNA (DOI: 10.5281/zenodo.8292325).

## ACKNOWLEDGMENTS

The authors thank Danny Incarnato for fruitful discussions. E.F.B. thanks his former supervisor Mikhail Roytberg for the visionary idea to choose a stem as the prediction unit, and Pavel Pevzner and Phillip Compeau for their online course “Bioinformatics Algorithms: An Active Learning Approach”, which had a huge impact on this work. E.F.B. was supported by the European Molecular Biology Organization [EMBO fellowship ALTF 525-2022 to E.F.B]. J.M.B. and G.I.N were supported by the Polish National Science Centre [NCN grant 2017/26/A/NZ1/01083 to J.M.B.].

## AUTHOR CONTRIBUTIONS

E.F.B. designed the computational method. E.F.B., D.R.B., and G.I.N. developed the computational tool. E.F.B. and D.R.B. performed the computational studies. All authors carried out the analysis and edited the manuscript. E.F.B. and J.M.B. supervised the work.

## COMPETING INTERESTS

The authors declare no competing interests.

## SUPPLEMENTARY MATERIALS

**Supplementary Figure S1.**
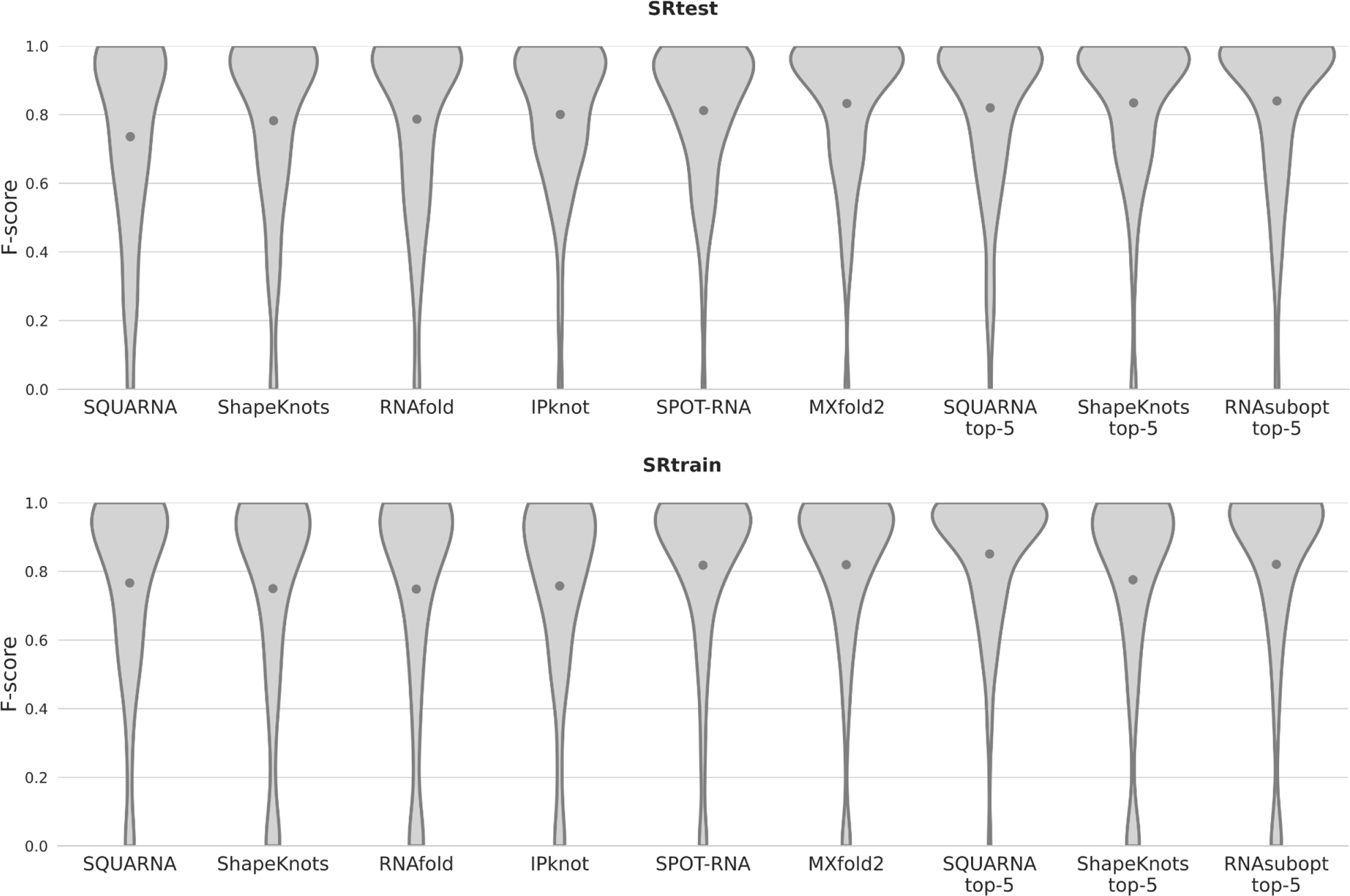
Single-sequence benchmark results. The violin plots illustrate F-score distributions derived from predictions of SQUARNA (top-1/top-5) and five state-of-the-art tools (RNAfold/RNAsubopt, IPknot, ShapeKnots, MXfold2, SPOT-RNA) on two RNA sequence datasets (SRtest with 247 sequences and SRtrain with 274 sequences). Mean values are indicated by gray dots. The graph was generated using the Seaborn Python library (https://seaborn.pydata.org/generated/seaborn.violinplot.html).

**Supplementary Figure S2.**
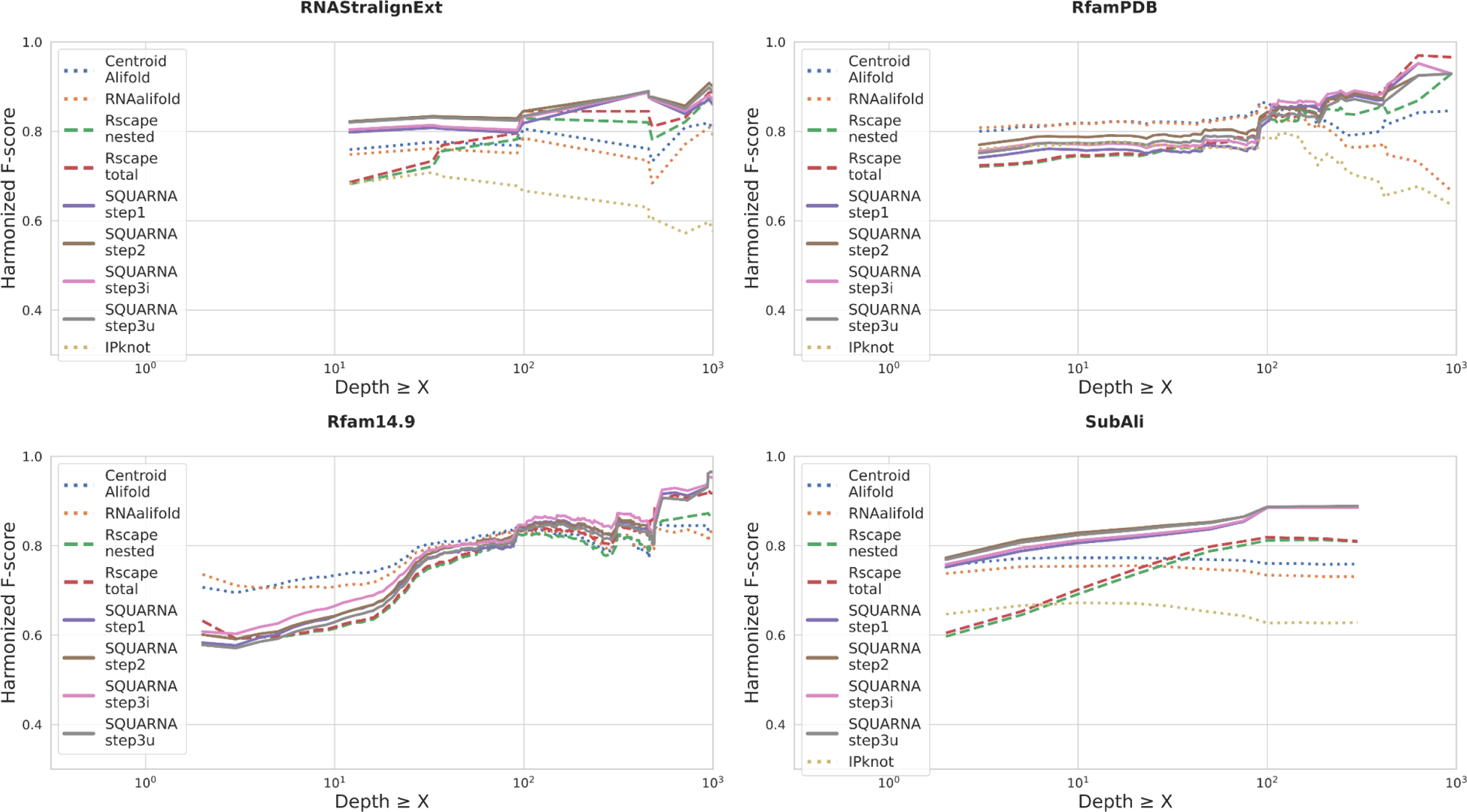
Benchmarking results for the alignment-based SQUARNA algorithm. X-axis - alignment depth threshold, Y-axis - harmonized F-score metric value. The four plots represent the four datasets: RNAStralignExt, RfamPDB, Rfam14.9, and SubAli.

**Supplementary Figure S3.**
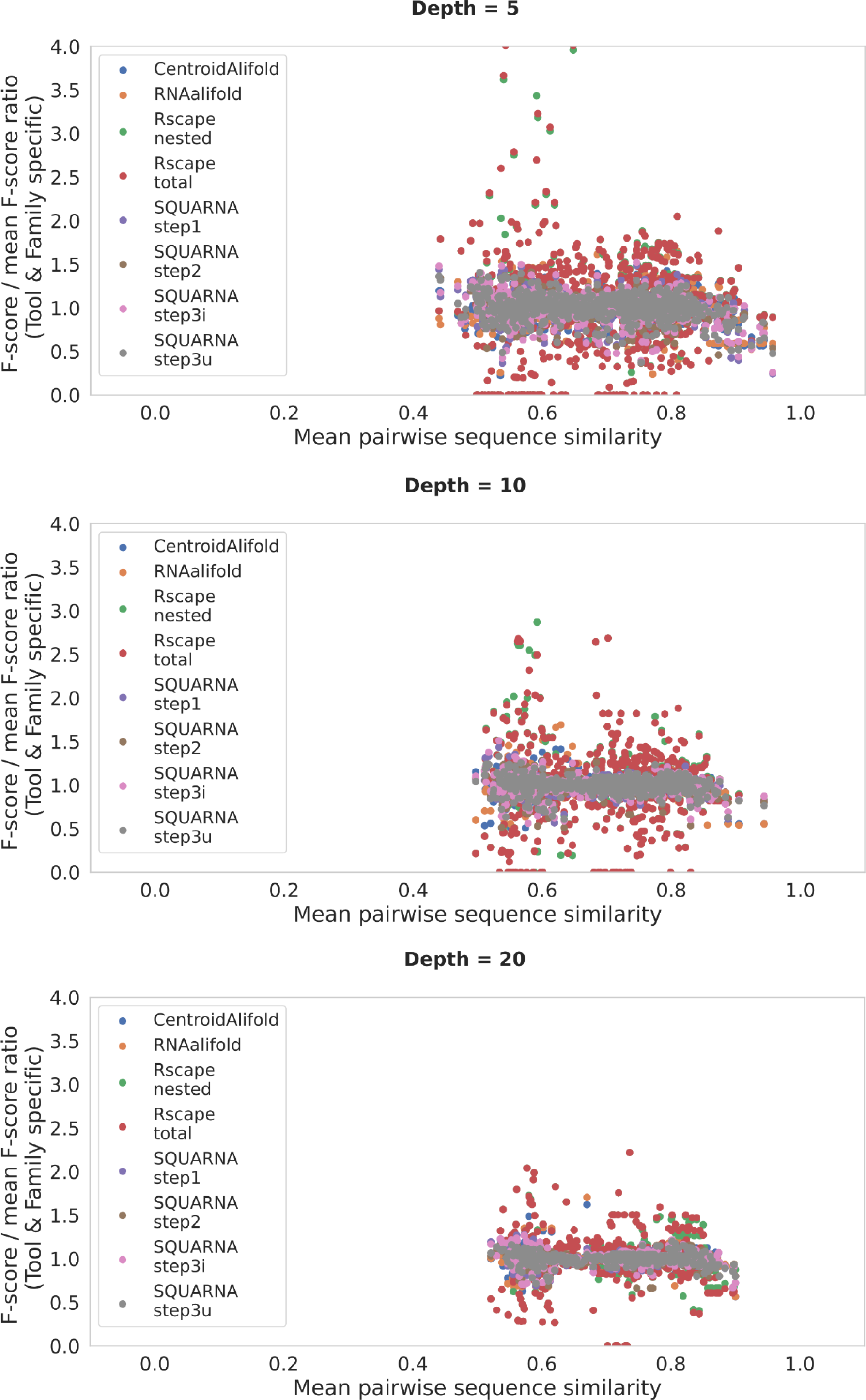
Dependence of the alignment-input tools’ performance on sequence similarity. The relative performance of alignment-input tools against the mean pairwise sequence similarity within the input alignment of depths 5, 10, and 20. The relative performance is measured as the ratio of an F-score to the mean F-score demonstrated by the tool on alignments of a specific family of the given depth. Rscape demonstrates the most unstable results across all presented alignment depths.

**Supplementary Figure S4.**
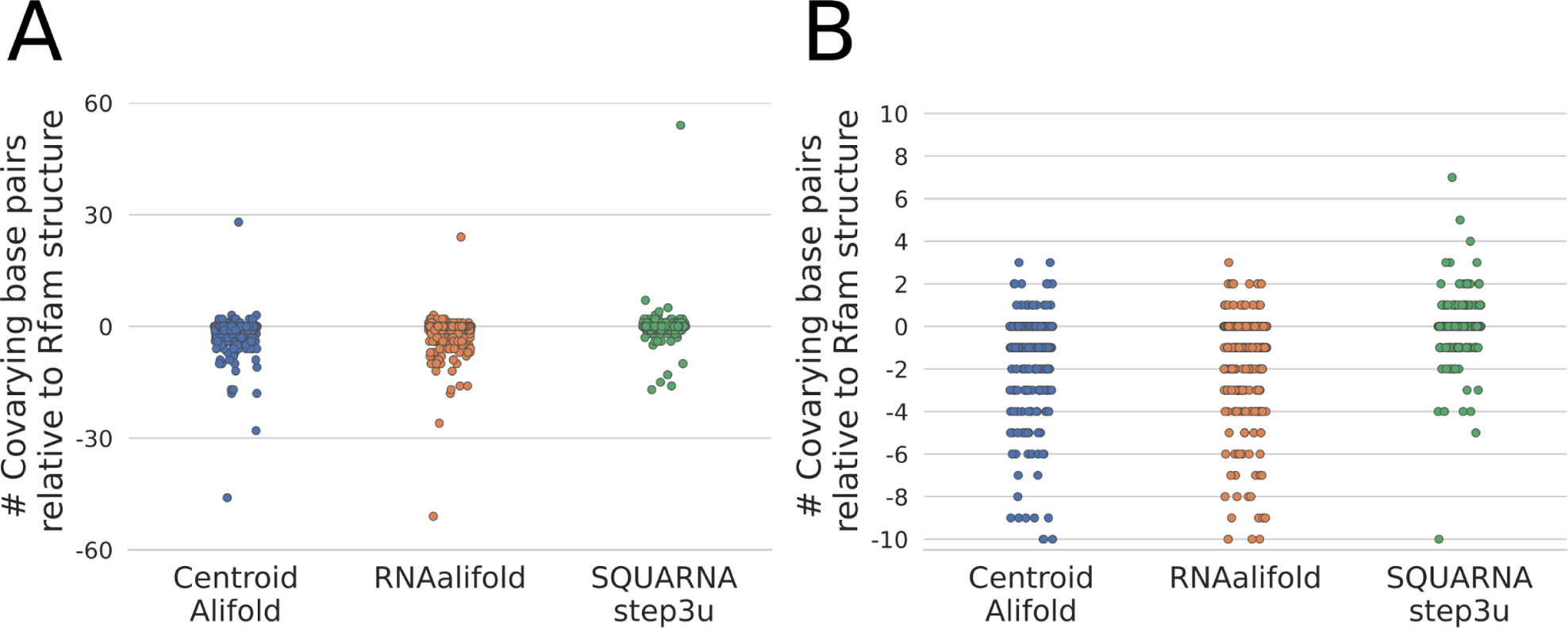
Benchmarking results in terms of covarying base pairs. Benchmarking results in the number of significantly covarying base pairs (measured by Rscape) among the base pairs predicted by CentroidAlifold, RNAalifold, and SQUARNA (SQUARNA step3u). Y axis shows the difference between the tool’s prediction and the Rfam annotated structure for 685 families of the Rfam14.9 dataset. Each dot represents a single Rfam family. (A) Outliers included; (B) Outliers excluded.

**Supplementary Figure S5.**
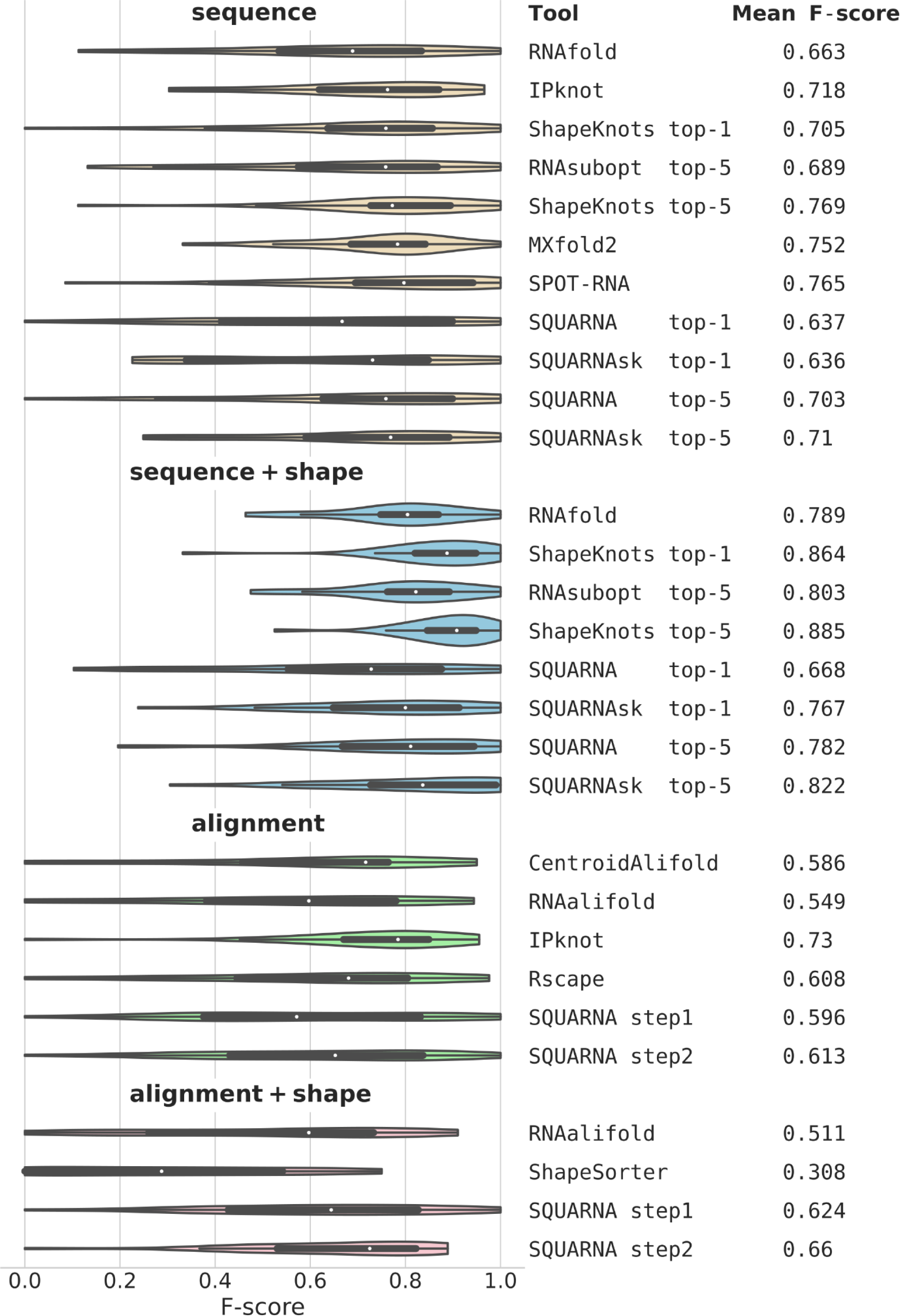
Benchmark results for the S01 dataset of 24 sequences. The violin plots represent F-score distributions resulting from predictions with four different input types: sequence, sequence with shape data, alignment, and alignment with shape data. Boxplots are enclosed within the violins, with white dots indicating median points. Mean F-scores are listed on the right side of the figure. The graph was prepared using the Seaborn Python library (https://seaborn.pydata.org/generated/seaborn.violinplot.html).

**Supplementary Figure S6.**
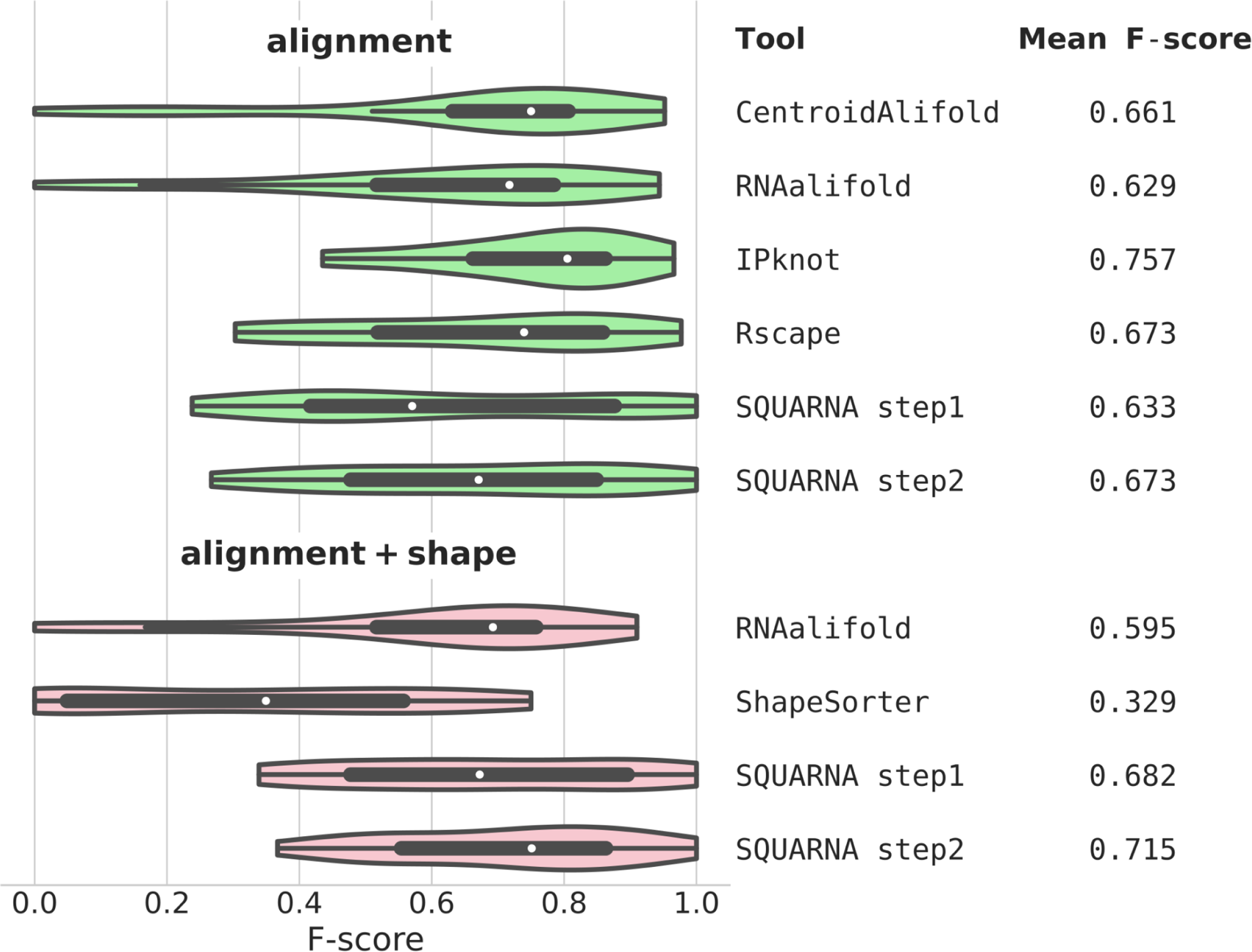
Benchmark results for the S01 dataset of 24 sequences. The violin plots represent F-score distributions resulting from predictions with two different input types: alignment only and alignment with shape data. The alignments were prepared in a structure-aware manner using covariation models. Boxplots are embedded within the violins, with white dots indicating median points. Mean F-scores are listed on the right side of the figure. The graph was prepared using the Seaborn Python library (https://seaborn.pydata.org/generated/seaborn.violinplot.html).

**Supplementary Figure S7.**
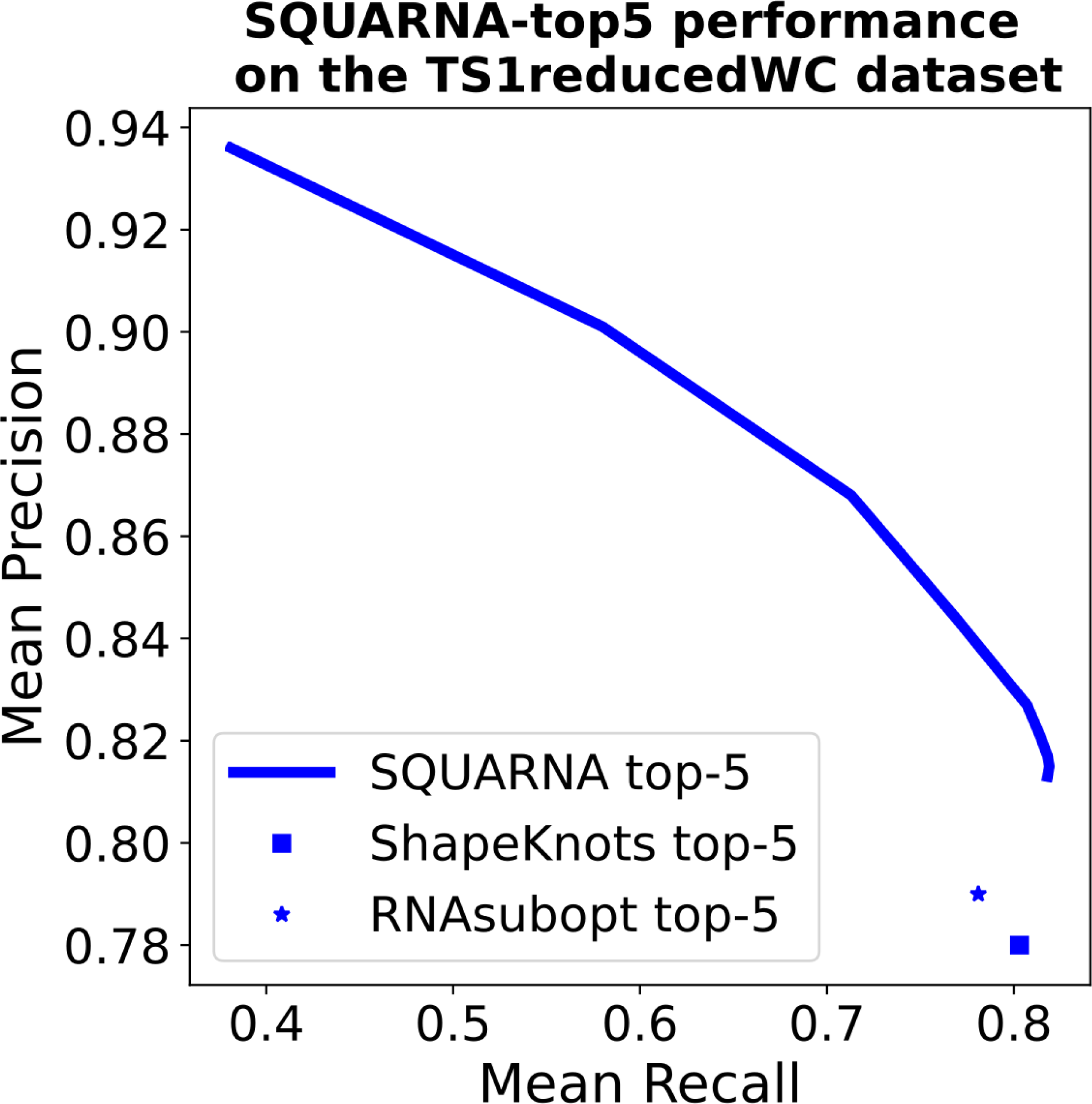
SQUARNA iterative prediction demonstration. The Precision-Recall curve obtained for the performance of SQUARNA-top5 in predicting from one (highest precision) to ten (lowest precision) stems for the sequences of the TS1reducedWC dataset.

**Supplementary Table S1.** Overview of the RNA secondary structure prediction tools used in this work. SQUARNA provides eight of the ten listed features, while the remaining tools offer a maximum of five features.

|  | RNAfold/<br>RNAsubopt | IPknot | Shape<br>Knots | MXfold2 | SPOT-RNA | SQUARNA | RNAalifold | CentroidAlifold | Rscape<br>(CaCoFold) |
| --- | --- | --- | --- | --- | --- | --- | --- | --- | --- |
| **single-sequence input** | **YES** | **YES** | **YES** | **YES** | **YES** | **YES** | NO | NO | NO |
| **alignment input** | NO | **YES** | NO | NO | NO | **YES** | **YES** | **YES** | **YES** |
| **structure restraints input** | **YES** | **YES** | **YES** | NO | NO | **YES** | **YES** | **YES** | NO |
| **residue reactivities input** | **YES** | NO | **YES** | NO | NO | **YES** | **YES** | NO | NO |
| **reference structure input** | NO | NO | NO | NO | NO | **YES** | NO | NO | **YES** |
| **circular molecule input** | **YES** | NO | NO | NO | NO | NO | **YES** | NO | NO |
| **multi-chain input** | NO | NO | NO | NO | NO | **YES** | NO | NO | NO |
| **predicts alternative structures** | **YES** | NO | **YES** | NO | NO | **YES** | NO | **YES** | NO |
| **predicts pseudoknots** | NO | **YES** | **YES** | NO | **YES** | **YES** | NO | NO | **YES** |
| **predicts non-canonical base pairs** | NO | NO | NO | NO | **YES** | NO | NO | **YES** | **YES** |
| **Number of features** | 5 | 4 | 5 | 1 | 3 | 8 | 4 | 4 | 4 |

**Supplementary Table S2.**
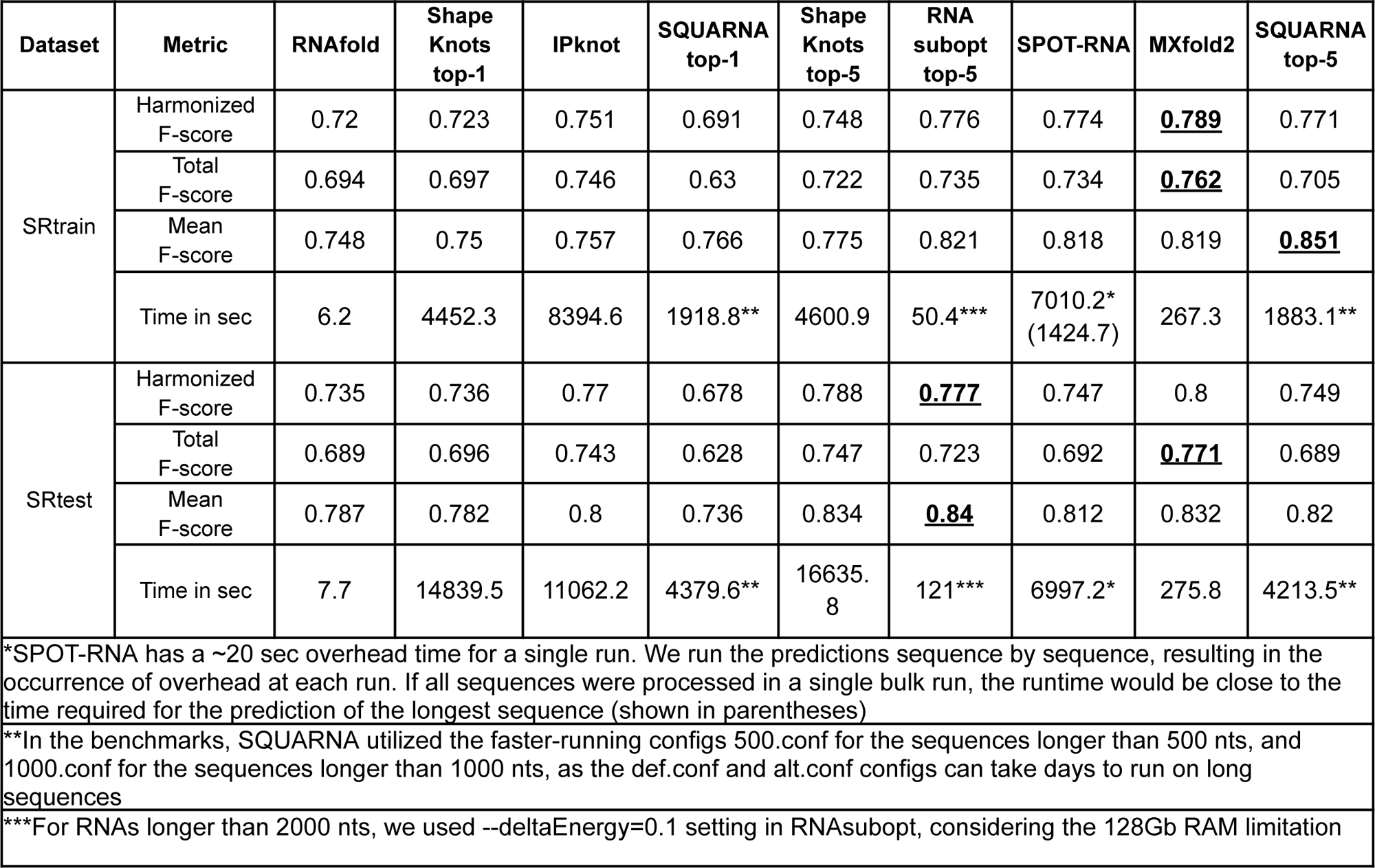
Comparisons of the single-sequence input tools.

**Supplementary Table S3.** Student’s paired t-test p-values calculated for each pair of the single-sequence input tools based on the sets of F-score values. Significant p-values (≤ 0.01) are highlighted in bold font.

| Pairwise t-test p-values on SRtrain dataset results |  |  |  |  |  |  |  |  |  |
| --- | --- | --- | --- | --- | --- | --- | --- | --- | --- |
| *SRtrain* | RNAfold | Shape<br>Knots<br>top-1 | IPknot | SQUARNA<br>top-1 | Shape<br>Knots<br>top-5 | RNA<br>subopt<br>top-5 | SPOT-RNA | MXfold2 | SQUARNA<br>top-5 |
| RNAfold | 1.00E+00 | 8.25E-01 | 5.47E-01 | 3.30E-01 | **7.41E-04** | **2.75E-10** | **5.78E-05** | **2.39E-05** | **1.59E-09** |
| Shape<br>Knots<br>top-1 | 8.25E-01 | 1.00E+00 | 6.03E-01 | 3.72E-01 | **2.07E-04** | **1.43E-08** | **8.50E-05** | **4.87E-05** | **2.80E-09** |
| IPknot | 5.47E-01 | 6.03E-01 | 1.00E+00 | 5.91E-01 | 2.11E-01 | **7.32E-06** | **4.95E-05** | **2.45E-04** | **1.35E-09** |
| SQUARNA<br>top-1 | 3.30E-01 | 3.72E-01 | 5.91E-01 | 1.00E+00 | 5.96E-01 | **6.15E-04** | **2.24E-03** | **9.96E-04** | **5.90E-13** |
| Shape<br>Knots<br>top-5 | **7.41E-04** | **2.07E-04** | 2.11E-01 | 5.96E-01 | 1.00E+00 | **1.95E-05** | **8.65E-03** | **7.41E-03** | **1.29E-06** |
| RNA<br>subopt<br>top-5 | **2.75E-10** | **1.43E-08** | **7.32E-06** | **6.15E-04** | **1.95E-05** | 1.00E+00 | 8.74E-01 | 9.47E-01 | 2.67E-02 |
| SPOT-RNA | **5.78E-05** | **8.50E-05** | **4.95E-05** | **2.24E-03** | **8.65E-03** | 8.74E-01 | 1.00E+00 | 9.13E-01 | 1.77E-02 |
| MXfold2 | **2.39E-05** | **4.87E-05** | **2.45E-04** | **9.96E-04** | **7.41E-03** | 9.47E-01 | 9.13E-01 | 1.00E+00 | 1.41E-02 |
| SQUARNA<br>top-5 | **1.59E-09** | **2.80E-09** | **1.35E-09** | **5.90E-13** | **1.29E-06** | 2.67E-02 | 1.77E-02 | 1.41E-02 | 1.00E+00 |
| Pairwise t-test p-values on SRtest dataset results |  |  |  |  |  |  |  |  |  |
| *SRtest* | RNAfold | Shape<br>Knots<br>top-1 | IPknot | SQUARNA<br>top-1 | Shape<br>Knots<br>top-5 | RNA<br>subopt<br>top-5 | SPOT-RNA | MXfold2 | SQUARNA<br>top-5 |
| RNAfold | 1.00E+00 | 7.08E-01 | 3.00E-01 | **2.37E-03** | **1.68E-04** | **1.14E-08** | 1.08E-01 | **8.19E-04** | 1.13E-02 |
| Shape<br>Knots<br>top-1 | 7.08E-01 | 1.00E+00 | 1.81E-01 | **9.73E-03** | **4.19E-08** | **1.74E-06** | 6.90E-02 | **6.78E-04** | 1.13E-02 |
| IPknot | 3.00E-01 | 1.81E-01 | 1.00E+00 | **7.76E-05** | **5.97E-03** | **2.09E-03** | 3.67E-01 | 2.16E-02 | 1.17E-01 |
| **SQUARNA**<br>**top-1** | **2.37E-03** | **9.73E-03** | **7.76E-05** | 1.00E+00 | **2.05E-08** | **7.50E-11** | **5.89E-07** | **7.37E-09** | **1.04E-13** |
| **Shape**<br>**Knots**<br>**top-5** | **1.68E-04** | **4.19E-08** | **5.97E-03** | **2.05E-08** | 1.00E+00 | 6.33E-01 | 1.03E-01 | 8.93E-01 | 2.74E-01 |
| **RNA**<br>**subopt**<br>**top-5** | **1.14E-08** | **1.74E-06** | **2.09E-03** | **7.50E-11** | 6.33E-01 | 1.00E+00 | 4.22E-02 | 5.17E-01 | 7.27E-02 |
| **SPOT-RNA** | 1.08E-01 | 6.90E-02 | 3.67E-01 | **5.89E-07** | 1.03E-01 | 4.22E-02 | 1.00E+00 | 1.01E-01 | 4.69E-01 |
| **MXfold2** | **8.19E-04** | **6.78E-04** | 2.16E-02 | **7.37E-09** | 8.93E-01 | 5.17E-01 | 1.01E-01 | 1.00E+00 | 3.35E-01 |
| **SQUARNA**<br>**top-5** | 1.13E-02 | 1.13E-02 | 1.17E-01 | **1.04E-13** | 2.74E-01 | 7.27E-02 | 4.69E-01 | 3.35E-01 | 1.00E+00 |

**Supplementary Table S4.** Contents of the RNAStralignExt dataset.

| RNAStralignExt |  |  |  |  |
| --- | --- | --- | --- | --- |
| N | Family | Rfam ID | Length | Depth |
| 1 | 5S rRNA | RF00001 | 230 | 712 |
| 2 | tRNA | RF00005 | 118 | 954 |
| 3 | RNaseP | RF00010 | 996 | 458 |
| 4 | tmRNA | RF00023 | 954 | 477 |
| 5 | telomerase RNA | RF00024 | 777 | 37 |
| 6 | group I intron | RF00028 | 892 | 12 |
| 7 | HDV ribozyme | RF00094 | 115 | 33 |
| 8 | SAM riboswitch | RF00162 | 239 | 457 |
| 9 | 16S rRNA | RF00177 | 1980 | 99 |
| 10 | SRP RNA | RF01854 | 315 | 92 |
| 11 | Twister ribozyme | RF03160 | 189 | 1613 |

**Supplementary Table S5.**
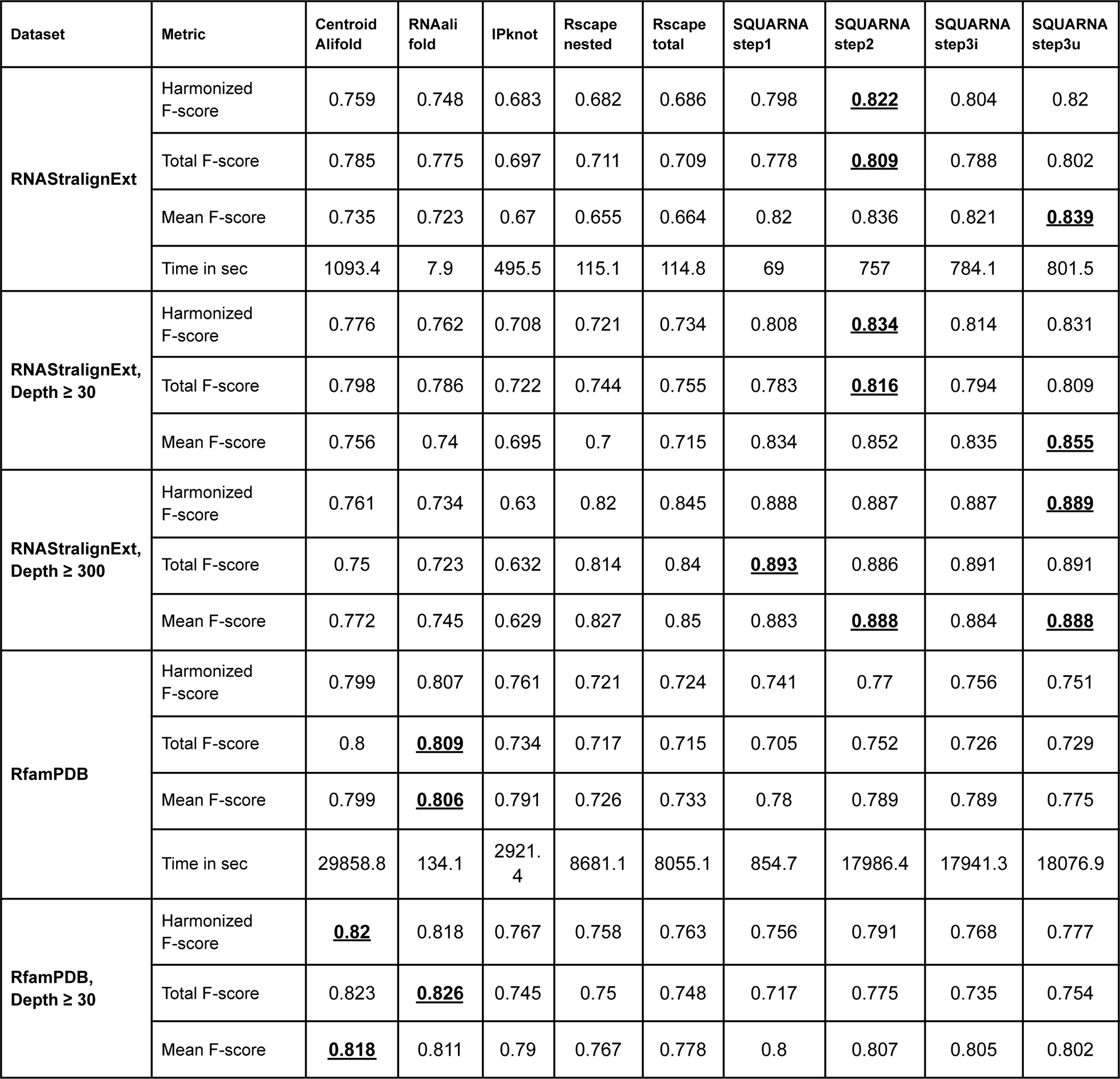
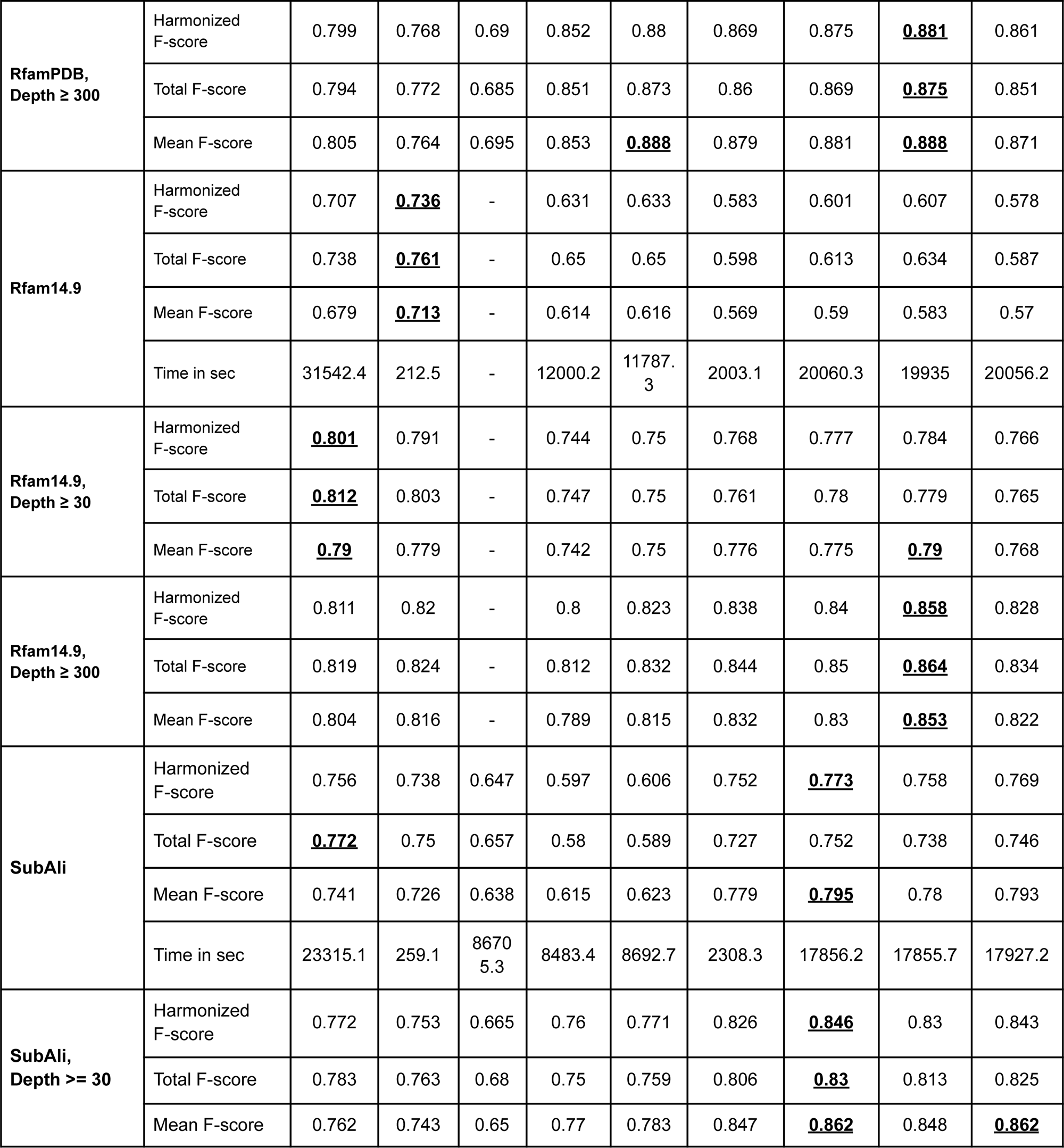
Comparisons of the alignment input tools. IPknot values are absent for the Rfam14.9 dataset as the execution of IPknot exceeded 10 days on the RF02746 family and was eventually aborted.

| Dataset | Metric | Centroid<br>Alifold | RNAali<br>fold | IPknot | Rscape<br>nested | Rscape<br>total | SQUARNA<br>step1 | SQUARNA<br>step2 | SQUARNA<br>step3i | SQUARNA<br>step3u |
| --- | --- | --- | --- | --- | --- | --- | --- | --- | --- | --- |
| RNAStralignExt | Harmonized F-score | 0.759 | 0.748 | 0.683 | 0.682 | 0.686 | 0.798 | <u>0.822</u> | 0.804 | 0.82 |
|  | Total F-score | 0.785 | 0.775 | 0.697 | 0.711 | 0.709 | 0.778 | <u>0.809</u> | 0.788 | 0.802 |
|  | Mean F-score | 0.735 | 0.723 | 0.67 | 0.655 | 0.664 | 0.82 | 0.836 | 0.821 | <u>0.839</u> |
|  | Time in sec | 1093.4 | 7.9 | 495.5 | 115.1 | 114.8 | 69 | 757 | 784.1 | 801.5 |
| RNAStralignExt,<br>Depth ≥ 30 | Harmonized F-score | 0.776 | 0.762 | 0.708 | 0.721 | 0.734 | 0.808 | <u>0.834</u> | 0.814 | 0.831 |
|  | Total F-score | 0.798 | 0.786 | 0.722 | 0.744 | 0.755 | 0.783 | <u>0.816</u> | 0.794 | 0.809 |
|  | Mean F-score | 0.756 | 0.74 | 0.695 | 0.7 | 0.715 | 0.834 | 0.852 | 0.835 | <u>0.855</u> |
| RNAStralignExt,<br>Depth ≥ 300 | Harmonized F-score | 0.761 | 0.734 | 0.63 | 0.82 | 0.845 | 0.888 | 0.887 | 0.887 | <u>0.889</u> |
|  | Total F-score | 0.75 | 0.723 | 0.632 | 0.814 | 0.84 | <u>0.893</u> | 0.886 | 0.891 | 0.891 |
|  | Mean F-score | 0.772 | 0.745 | 0.629 | 0.827 | 0.85 | 0.883 | <u>0.888</u> | 0.884 | <u>0.888</u> |
| RfamPDB | Harmonized F-score | 0.799 | 0.807 | 0.761 | 0.721 | 0.724 | 0.741 | 0.77 | 0.756 | 0.751 |
|  | Total F-score | 0.8 | <u>0.809</u> | 0.734 | 0.717 | 0.715 | 0.705 | 0.752 | 0.726 | 0.729 |
|  | Mean F-score | 0.799 | <u>0.806</u> | 0.791 | 0.726 | 0.733 | 0.78 | 0.789 | 0.789 | 0.775 |
|  | Time in sec | 29858.8 | 134.1 | 2921.4 | 8681.1 | 8055.1 | 854.7 | 17986.4 | 17941.3 | 18076.9 |
| RfamPDB,<br>Depth ≥ 30 | Harmonized F-score | <u>0.82</u> | 0.818 | 0.767 | 0.758 | 0.763 | 0.756 | 0.791 | 0.768 | 0.777 |
|  | Total F-score | 0.823 | <u>0.826</u> | 0.745 | 0.75 | 0.748 | 0.717 | 0.775 | 0.735 | 0.754 |
|  | Mean F-score | <u>0.818</u> | 0.811 | 0.79 | 0.767 | 0.778 | 0.8 | 0.807 | 0.805 | 0.802 |
| **RfamPDB,**<br>**Depth ≥ 300** | Harmonized F-score | 0.799 | 0.768 | 0.69 | 0.852 | 0.88 | 0.869 | 0.875 | **<u>0.881</u>** | 0.861 |
|  | Total F-score | 0.794 | 0.772 | 0.685 | 0.851 | 0.873 | 0.86 | 0.869 | **<u>0.875</u>** | 0.851 |
|  | Mean F-score | 0.805 | 0.764 | 0.695 | 0.853 | **<u>0.888</u>** | 0.879 | 0.881 | **<u>0.888</u>** | 0.871 |
| **Rfam14.9** | Harmonized F-score | 0.707 | **<u>0.736</u>** | - | 0.631 | 0.633 | 0.583 | 0.601 | 0.607 | 0.578 |
|  | Total F-score | 0.738 | **<u>0.761</u>** | - | 0.65 | 0.65 | 0.598 | 0.613 | 0.634 | 0.587 |
|  | Mean F-score | 0.679 | **<u>0.713</u>** | - | 0.614 | 0.616 | 0.569 | 0.59 | 0.583 | 0.57 |
|  | Time in sec | 31542.4 | 212.5 | - | 12000.2 | 11787.3 | 2003.1 | 20060.3 | 19935 | 20056.2 |
| **Rfam14.9,**<br>**Depth ≥ 30** | Harmonized F-score | **<u>0.801</u>** | 0.791 | - | 0.744 | 0.75 | 0.768 | 0.777 | 0.784 | 0.766 |
|  | Total F-score | **<u>0.812</u>** | 0.803 | - | 0.747 | 0.75 | 0.761 | 0.78 | 0.779 | 0.765 |
|  | Mean F-score | **<u>0.79</u>** | 0.779 | - | 0.742 | 0.75 | 0.776 | 0.775 | **<u>0.79</u>** | 0.768 |
| **Rfam14.9,**<br>**Depth ≥ 300** | Harmonized F-score | 0.811 | 0.82 | - | 0.8 | 0.823 | 0.838 | 0.84 | **<u>0.858</u>** | 0.828 |
|  | Total F-score | 0.819 | 0.824 | - | 0.812 | 0.832 | 0.844 | 0.85 | **<u>0.864</u>** | 0.834 |
|  | Mean F-score | 0.804 | 0.816 | - | 0.789 | 0.815 | 0.832 | 0.83 | **<u>0.853</u>** | 0.822 |
| **SubAli** | Harmonized F-score | 0.756 | 0.738 | 0.647 | 0.597 | 0.606 | 0.752 | **<u>0.773</u>** | 0.758 | 0.769 |
|  | Total F-score | **<u>0.772</u>** | 0.75 | 0.657 | 0.58 | 0.589 | 0.727 | 0.752 | 0.738 | 0.746 |
|  | Mean F-score | 0.741 | 0.726 | 0.638 | 0.615 | 0.623 | 0.779 | **<u>0.795</u>** | 0.78 | 0.793 |
|  | Time in sec | 23315.1 | 259.1 | 86705.3 | 8483.4 | 8692.7 | 2308.3 | 17856.2 | 17855.7 | 17927.2 |
| **SubAli,**<br>**Depth ≥ 30** | Harmonized F-score | 0.772 | 0.753 | 0.665 | 0.76 | 0.771 | 0.826 | **<u>0.846</u>** | 0.83 | 0.843 |
|  | Total F-score | 0.783 | 0.763 | 0.68 | 0.75 | 0.759 | 0.806 | **<u>0.83</u>** | 0.813 | 0.825 |
|  | Mean F-score | 0.762 | 0.743 | 0.65 | 0.77 | 0.783 | 0.847 | **<u>0.862</u>** | 0.848 | **<u>0.862</u>** |

**Supplementary Table S6.** Student’s paired t-test p-values calculated for each pair of the alignment input tools based on the sets of F-score values. Significant p-values (≤ 0.01) are highlighted in bold font.

**Paired t-test p-values on SubAli dataset results**
| SubAli | CentroidAlifold | RNAalifold | IPknot | Rscape nested | Rscape total | SQUARNA step1 | SQUARNA step2 | SQUARNA step3i | SQUARNA step3u |
| --- | --- | --- | --- | --- | --- | --- | --- | --- | --- |
| CentroidAlifold | 1.00E+00 | **<u>5.24E-17</u>** | **<u>1.82E-76</u>** | **<u>7.05E-56</u>** | **<u>3.48E-46</u>** | **<u>1.14E-14</u>** | **<u>4.58E-29</u>** | **<u>2.18E-15</u>** | **<u>1.33E-27</u>** |
| RNAalifold | **<u>5.24E-17</u>** | 1.00E+00 | **<u>1.27E-53</u>** | **<u>5.91E-42</u>** | **<u>2.92E-34</u>** | **<u>2.55E-25</u>** | **<u>1.03E-41</u>** | **<u>1.23E-26</u>** | **<u>1.58E-39</u>** |
| IPknot | **<u>1.82E-76</u>** | **<u>1.27E-53</u>** | 1.00E+00 | 1.10E-02 | 1.06E-01 | **<u>1.71E-100</u>** | **<u>4.23E-126</u>** | **<u>9.31E-97</u>** | **<u>2.11E-128</u>** |
| Rscape nested | **<u>7.05E-56</u>** | **<u>5.91E-42</u>** | 1.10E-02 | 1.00E+00 | **<u>1.16E-08</u>** | **<u>7.11E-97</u>** | **<u>6.36E-113</u>** | **<u>1.99E-95</u>** | **<u>1.82E-113</u>** |
| Rscape total | **<u>3.48E-46</u>** | **<u>2.92E-34</u>** | 1.06E-01 | **<u>1.16E-08</u>** | 1.00E+00 | **<u>5.74E-88</u>** | **<u>4.19E-104</u>** | **<u>8.87E-87</u>** | **<u>1.89E-104</u>** |
| SQUARNA step1 | **<u>1.14E-14</u>** | **<u>2.55E-25</u>** | **<u>1.71E-100</u>** | **<u>7.11E-97</u>** | **<u>5.74E-88</u>** | 1.00E+00 | **<u>5.00E-14</u>** | 2.75E-01 | **<u>9.40E-22</u>** |
| SQUARNA step2 | **<u>4.58E-29</u>** | **<u>1.03E-41</u>** | **<u>4.23E-126</u>** | **<u>6.36E-113</u>** | **<u>4.19E-104</u>** | **<u>5.00E-14</u>** | 1.00E+00 | **<u>1.12E-17</u>** | 2.43E-01 |
| SQUARNA step3i | **<u>2.18E-15</u>** | **<u>1.23E-26</u>** | **<u>9.31E-97</u>** | **<u>1.99E-95</u>** | **<u>8.87E-87</u>** | 2.75E-01 | **<u>1.12E-17</u>** | 1.00E+00 | **<u>1.67E-12</u>** |
| SQUARNA step3u | **<u>1.33E-27</u>** | **<u>1.58E-39</u>** | **<u>2.11E-128</u>** | **<u>1.82E-113</u>** | **<u>1.89E-104</u>** | **<u>9.40E-22</u>** | 2.43E-01 | **<u>1.67E-12</u>** | 1.00E+00 |

**Paired t-test p-values on SubAli, Depth $\geq 30$ dataset results**
| SubAli,<br>Depth $\geq 30$ | CentroidAlifold | RNAalifold | IPknot | Rscape nested | Rscape total | SQUARNA step1 | SQUARNA step2 | SQUARNA step3i | SQUARNA step3u |
| --- | --- | --- | --- | --- | --- | --- | --- | --- | --- |
| CentroidAlifold | 1.00E+00 | **<u>1.21E-42</u>** | **<u>9.79E-46</u>** | 1.21E-01 | **<u>6.76E-04</u>** | **<u>9.37E-44</u>** | **<u>7.50E-61</u>** | **<u>9.70E-48</u>** | **<u>2.00E-56</u>** |
| RNAalifold | **<u>1.21E-42</u>** | 1.00E+00 | **<u>7.58E-33</u>** | **<u>4.24E-07</u>** | **<u>2.49E-10</u>** | **<u>1.23E-57</u>** | **<u>2.80E-76</u>** | **<u>3.42E-62</u>** | **<u>2.52E-71</u>** |
| IPknot | **<u>9.79E-46</u>** | **<u>7.58E-33</u>** | 1.00E+00 | **<u>2.43E-33</u>** | **<u>1.50E-36</u>** | **<u>4.89E-119</u>** | **<u>2.68E-141</u>** | **<u>2.76E-117</u>** | **<u>6.04E-143</u>** |
| Rscape nested | 1.21E-01 | **<u>4.24E-07</u>** | **<u>2.43E-33</u>** | 1.00E+00 | **<u>1.64E-07</u>** | **<u>3.05E-36</u>** | **<u>3.48E-49</u>** | **<u>6.11E-38</u>** | **<u>6.01E-47</u>** |
| Rscape<br>total | <u>6.76E-04</u> | <u>2.49E-10</u> | <u>1.50E-36</u> | <u>1.64E-07</u> | 1.00E+00 | <u>5.01E-27</u> | <u>3.07E-40</u> | <u>1.50E-28</u> | <u>4.91E-38</u> |
| SQUARNA<br>step1 | <u>9.37E-44</u> | <u>1.23E-57</u> | <u>4.89E-119</u> | <u>3.05E-36</u> | <u>5.01E-27</u> | 1.00E+00 | <u>2.33E-15</u> | 2.94E-01 | <u>7.13E-18</u> |
| SQUARNA<br>step2 | <u>7.50E-61</u> | <u>2.80E-76</u> | <u>2.68E-141</u> | <u>3.48E-49</u> | <u>3.07E-40</u> | <u>2.33E-15</u> | 1.00E+00 | <u>1.16E-14</u> | 8.15E-01 |
| SQUARNA<br>step3i | <u>9.70E-48</u> | <u>3.42E-62</u> | <u>2.76E-117</u> | <u>6.11E-38</u> | <u>1.50E-28</u> | 2.94E-01 | <u>1.16E-14</u> | 1.00E+00 | <u>2.51E-12</u> |
| SQUARNA<br>step3u | <u>2.00E-56</u> | <u>2.52E-71</u> | <u>6.04E-143</u> | <u>6.01E-47</u> | <u>4.91E-38</u> | <u>7.13E-18</u> | 8.15E-01 | <u>2.51E-12</u> | 1.00E+00 |

**Supplementary Table S7.**
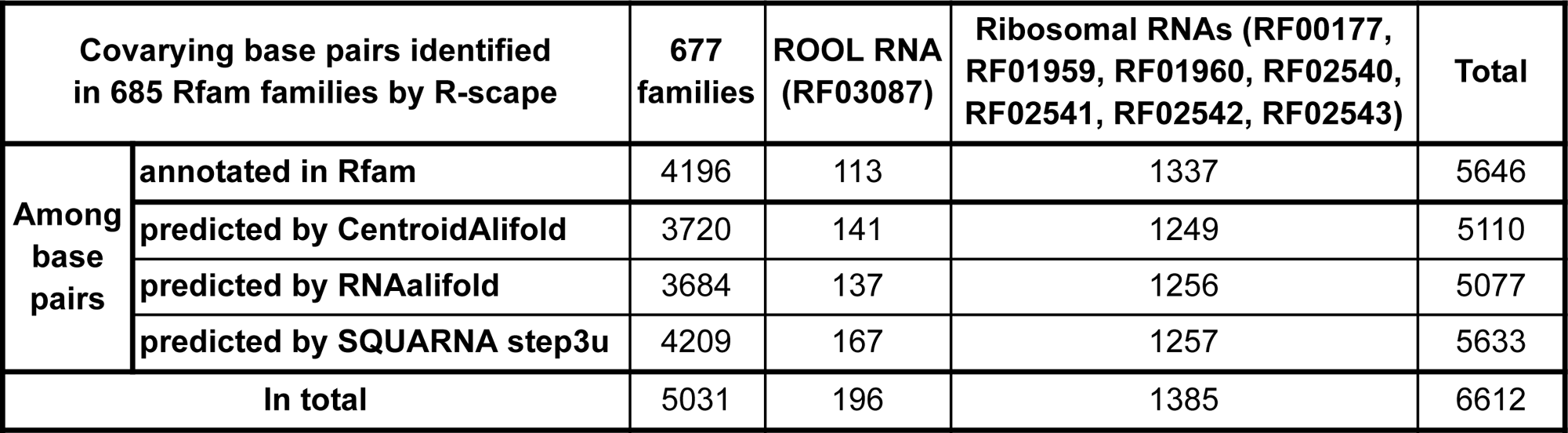
Comparisons in terms of covarying base pairs as measured by Rscape.

| Covarying base pairs identified in 685 Rfam families by R-scape |  | 677 families | ROOL RNA (RF03087) | Ribosomal RNAs (RF00177, RF01959, RF01960, RF02540, RF02541, RF02542, RF02543) | Total |
| --- | --- | --- | --- | --- | --- |
| Among base pairs | annotated in Rfam | 4196 | 113 | 1337 | 5646 |
|  | predicted by CentroidAlifold | 3720 | 141 | 1249 | 5110 |
|  | predicted by RNAalifold | 3684 | 137 | 1256 | 5077 |
|  | predicted by SQUARNA step3u | 4209 | 167 | 1257 | 5633 |
| In total |  | 5031 | 196 | 1385 | 6612 |

**Supplementary Table S8.** Comparisons of the RNA secondary structure prediction tools on the S01 dataset with different input types.

| Metrics type |  | Harmonic (Mean, Total) |  |  | Total |  |  | Mean |  |  | Time in sec |
| --- | --- | --- | --- | --- | --- | --- | --- | --- | --- | --- | --- |
| Metrics |  | F-score | Precision | Recall | F-score | Precision | Recall | F-score | Precision | Recall |  |
| Input | Tool |  |  |  |  |  |  |  |  |  |  |
| sequence | RNAfold | 0.639 | 0.647 | 0.637 | 0.617 | 0.609 | 0.626 | 0.663 | 0.689 | 0.648 | 0.9 |
|  | RNAsubopt top-5 | 0.666 | 0.672 | 0.664 | 0.644 | 0.635 | 0.653 | 0.689 | 0.714 | 0.676 | 1.9 |
|  | IPknot | 0.701 | 0.766 | 0.65 | 0.684 | 0.745 | 0.633 | 0.718 | 0.788 | 0.668 | 17.7 |
|  | ShapeKnots top-1 | 0.681 | 0.678 | 0.688 | 0.659 | 0.645 | 0.674 | 0.705 | 0.714 | 0.702 | 287.5 |
|  | ShapeKnots top-5 | <u>0.749</u> | 0.746 | <u>0.754</u> | 0.73 | 0.718 | <u>0.743</u> | <u>0.769</u> | 0.777 | <u>0.766</u> | 278.9 |
|  | MXfold2 | 0.744 | 0.774 | 0.721 | <u>0.736</u> | 0.749 | 0.724 | 0.752 | 0.801 | 0.719 | 26.4 |
|  | SPOT-RNA | 0.719 | <u>0.837</u> | 0.638 | 0.678 | <u>0.818</u> | 0.579 | 0.765 | <u>0.856</u> | 0.71 | 852.7*<br>(72.7) |
|  | SQUARNA top-1 | 0.602 | 0.613 | 0.595 | 0.571 | 0.576 | 0.566 | 0.637 | 0.655 | 0.628 | 1032.5** |
|  | SQUARNAsk top-1 | 0.605 | 0.603 | 0.609 | 0.576 | 0.567 | 0.585 | 0.636 | 0.644 | 0.636 | 136.3** |
|  | SQUARNA top-5 | 0.658 | 0.677 | 0.644 | 0.618 | 0.628 | 0.609 | 0.703 | 0.734 | 0.683 | 1016.6** |
|  | SQUARNAsk top-5 | 0.666 | 0.676 | 0.661 | 0.627 | 0.628 | 0.627 | 0.71 | 0.733 | 0.699 | 136.4** |
| sequence + shape | RNAfold | 0.778 | 0.802 | 0.76 | 0.768 | 0.78 | 0.756 | 0.789 | 0.825 | 0.764 | 1.2 |
|  | RNAsubopt top-5 | 0.792 | 0.813 | 0.774 | 0.781 | 0.791 | 0.771 | 0.803 | 0.837 | 0.778 | 1.9 |
|  | ShapeKnots top-1 | 0.852 | 0.844 | 0.862 | 0.841 | 0.831 | 0.852 | 0.864 | 0.858 | 0.872 | 292.6 |
|  | ShapeKnots top-5 | <u>0.873</u> | <u>0.864</u> | <u>0.883</u> | <u>0.862</u> | <u>0.851</u> | <u>0.874</u> | <u>0.885</u> | <u>0.878</u> | <u>0.893</u> | 318.1 |
|  | SQUARNA top-1 | 0.649 | 0.635 | 0.666 | 0.631 | 0.618 | 0.646 | 0.668 | 0.653 | 0.687 | 1927.8** |
|  | SQUARNAsk top-1 | 0.737 | 0.725 | 0.75 | 0.709 | 0.698 | 0.72 | 0.767 | 0.755 | 0.783 | 214.4** |
|  | SQUARNA top-5 | 0.734 | 0.72 | 0.749 | 0.691 | 0.678 | 0.705 | 0.782 | 0.768 | 0.799 | 1914.7** |
|  | SQUARNAsk top-5 | 0.779 | 0.778 | 0.781 | 0.74 | 0.737 | 0.743 | 0.822 | 0.823 | 0.824 | 212.8** |
| alignment | CentroidAlifold | 0.606 | 0.841 | 0.496 | 0.627 | <u>0.855</u> | 0.494 | 0.586 | 0.828 | 0.498 | 239.8 |
|  | RNAalifold | 0.565 | 0.729 | 0.489 | 0.583 | 0.717 | 0.491 | 0.549 | 0.741 | 0.488 | 8 |
|  | IPknot | <u>0.719</u> | <u>0.835</u> | <u>0.636</u> | <u>0.709</u> | 0.827 | <u>0.62</u> | <u>0.73</u> | <u>0.844</u> | <u>0.653</u> | 239.8 |
|  | Rscape nested | 0.589 | 0.652 | 0.542 | 0.571 | 0.62 | 0.529 | 0.608 | 0.687 | 0.555 | 89.8 |
|  | Rscape total | 0.589 | 0.652 | 0.542 | 0.571 | 0.62 | 0.529 | 0.608 | 0.687 | 0.555 | 89.2 |
|  | SQUARNA step1 | 0.553 | 0.627 | 0.5 | 0.515 | 0.606 | 0.448 | 0.596 | 0.65 | 0.565 | 26.9 |
|  | SQUARNA step2 | 0.578 | 0.587 | 0.573 | 0.547 | 0.563 | 0.532 | 0.613 | 0.614 | 0.622 | 129 |
|  | SQUARNA step3i | 0.577 | 0.767 | 0.478 | 0.545 | 0.763 | 0.424 | 0.612 | 0.772 | 0.549 | 129.9 |
|  | SQUARNA step3u | 0.555 | 0.557 | 0.555 | 0.52 | 0.53 | 0.51 | 0.594 | 0.587 | 0.609 | 130.2 |
| alignment<br>+ shape | RNAalifold | 0.528 | 0.696 | 0.454 | 0.547 | 0.675 | 0.46 | 0.511 | 0.719 | 0.449 | 9.5 |
|  | ShapeSorter*** | 0.316 | <u>0.863</u> | 0.208 | 0.325 | <u>0.841</u> | 0.201 | 0.308 | <u>0.887</u> | 0.216 | _<sup>***</sup> |
|  | SQUARNA step1 | 0.591 | 0.649 | 0.547 | 0.562 | 0.638 | 0.502 | 0.624 | 0.66 | 0.601 | 35.8 |
|  | SQUARNA step2 | <u>0.639</u> | 0.637 | <u>0.646</u> | <u>0.62</u> | 0.629 | <u>0.612</u> | <u>0.66</u> | 0.646 | <u>0.685</u> | 142.2 |
|  | SQUARNA step3i | 0.629 | 0.788 | 0.533 | 0.602 | 0.789 | 0.486 | 0.659 | 0.787 | 0.59 | 143.2 |
|  | SQUARNA step3u | 0.603 | 0.598 | 0.611 | 0.58 | 0.586 | 0.574 | 0.627 | 0.61 | 0.653 | 140.2 |

\*SPOT-RNA has a ~20 sec overhead time for a single run. We run the predictions sequence by sequence, resulting in the occurrence of overhead at each run. If all sequences were processed in a single bulk run, the runtime would be close to the time required for the prediction of the longest sequence (shown in parentheses)
**\*\***In the benchmarks, SQUARNA utilized the faster-running configs 500.conf for the sequences longer than 500 nts, and 1000.conf for the sequences longer than 1000 nts, as the def.conf and alt.conf configs can take days to run on long sequences.
\*\*\*We utilized the ShapeSorter web server, and hence, the running time could not be measured.

**Supplementary Table S9.**
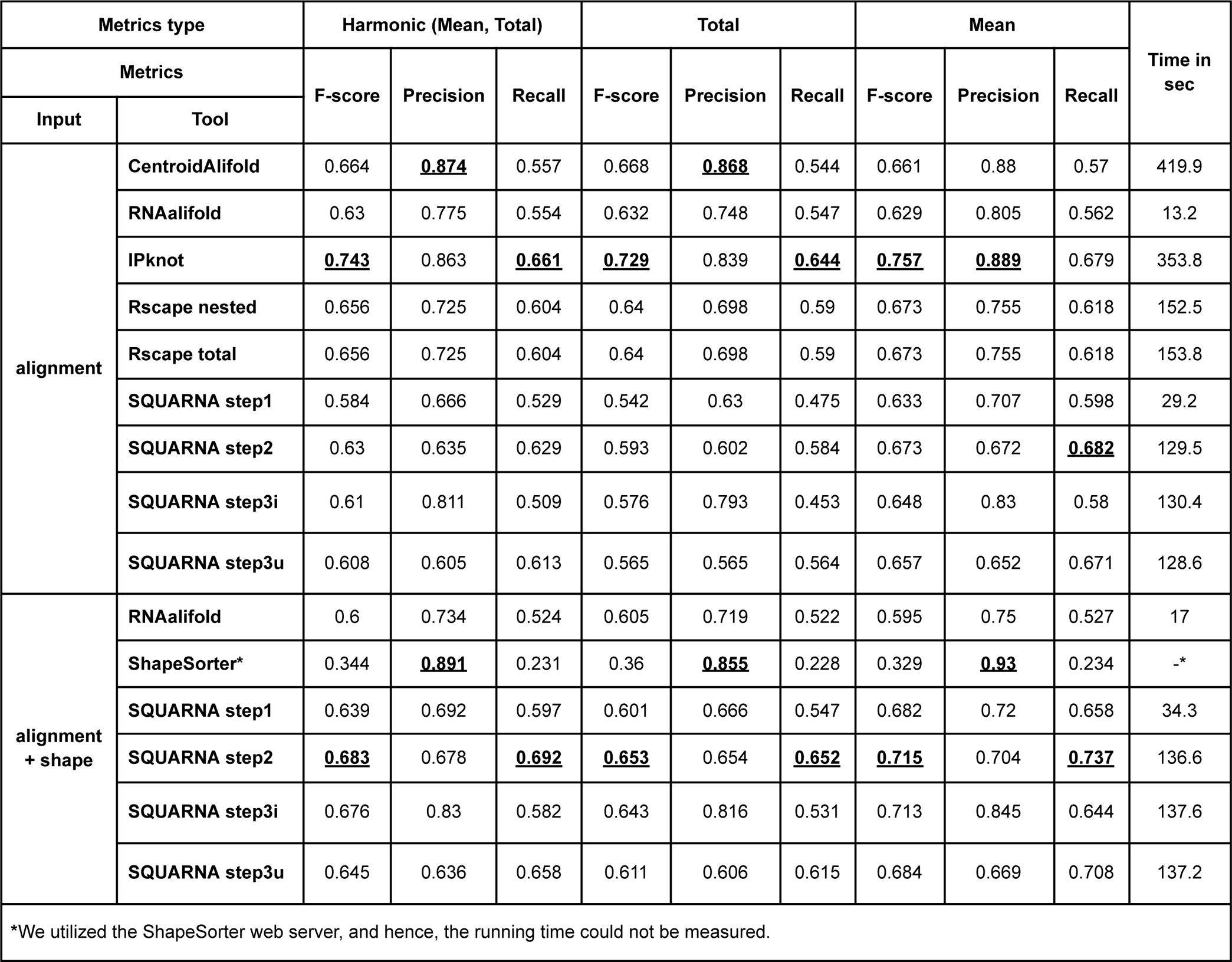
Comparisons of the RNA secondary structure prediction tools on the S01 dataset with structural alignment input types built with covariation models.

**Supplementary Table S10.** Comparisons of the RNA secondary structure prediction tools on the Ribonanza dataset with sequence-only input and with sequence plus DMS/2A3 reactivitites input data.

| Metrics type |  | Harmonic (Mean, Total) |  |  | Total |  |  | Mean |  |  | Time in sec |
| --- | --- | --- | --- | --- | --- | --- | --- | --- | --- | --- | --- |
| Metrics |  | F-score | Precision | Recall | F-score | Precision | Recall | F-score | Precision | Recall |  |
| Input | Tool |  |  |  |  |  |  |  |  |  |  |
| sequence | RNAfold | 0.688 | 0.741 | 0.652 | 0.709 | 0.744 | 0.677 | 0.668 | 0.739 | 0.629 | 0.4 |
|  | RNAsubopt top-5 | 0.711 | 0.784 | 0.663 | 0.73 | 0.778 | 0.688 | 0.693 | 0.79 | 0.64 | 0.5 |
|  | IPknot | 0.794 | 0.826 | 0.773 | 0.797 | 0.817 | 0.778 | 0.792 | 0.836 | 0.768 | 1 |
|  | ShapeKnots top-1 | 0.757 | 0.766 | 0.752 | 0.758 | 0.762 | 0.754 | 0.756 | 0.77 | 0.75 | 1590.8 |
|  | ShapeKnots top-5 | <u>0.839</u> | 0.843 | <u>0.839</u> | <u>0.847</u> | 0.845 | <u>0.849</u> | 0.831 | 0.841 | <u>0.83</u> | 1599.6 |
|  | MXfold2 | 0.692 | 0.819 | 0.625 | 0.716 | 0.812 | 0.64 | 0.669 | 0.826 | 0.61 | 60.3 |
|  | SPOT-RNA | 0.828 | 0.873 | 0.796 | 0.835 | 0.878 | 0.796 | 0.822 | 0.869 | 0.796 | 655.3*<br>(32.2) |
|  | SQUARNA top-1 | 0.765 | 0.786 | 0.749 | 0.781 | 0.808 | 0.757 | 0.75 | 0.765 | <u>0.741</u> | 7.9 |
|  | SQUARNAalt top-1 | 0.81 | 0.826 | 0.797 | 0.823 | 0.842 | 0.804 | 0.797 | 0.81 | <u>0.79</u> | 6 |
|  | SQUARNAsk top-1 | 0.738 | 0.764 | 0.718 | 0.751 | 0.782 | 0.722 | 0.726 | 0.746 | <u>0.715</u> | 7.5 |
|  | SQUARNA top-5 | 0.833 | 0.882 | 0.799 | 0.84 | 0.883 | 0.802 | 0.827 | 0.881 | 0.796 | 7.9 |
|  | SQUARNAalt top-5 | <u>0.839</u> | 0.862 | 0.824 | 0.846 | 0.865 | 0.828 | <u>0.832</u> | 0.859 | 0.82 | 5.2 |
|  | SQUARNAsk top-5 | 0.82 | <u>0.889</u> | 0.772 | 0.827 | <u>0.893</u> | 0.77 | 0.814 | <u>0.885</u> | 0.774 | 7.5 |
| sequence + DMS | RNAfold | 0.646 | 0.714 | 0.604 | 0.673 | 0.72 | 0.632 | 0.621 | 0.708 | 0.579 | 0.3 |
|  | RNAsubopt top-5 | 0.655 | 0.729 | 0.608 | 0.682 | 0.736 | 0.635 | 0.63 | 0.722 | 0.583 | 0.8 |
|  | ShapeKnots top-1 | 0.725 | 0.734 | 0.724 | 0.74 | 0.747 | 0.733 | 0.711 | 0.722 | 0.715 | 1785.7 |
|  | ShapeKnots top-5 | 0.797 | 0.812 | 0.792 | 0.813 | 0.822 | 0.804 | 0.782 | 0.802 | 0.781 | 1802.3 |
|  | SQUARNA top-1 | 0.814 | 0.814 | 0.815 | 0.818 | 0.821 | 0.815 | 0.81 | 0.808 | 0.815 | 10 |
|  | SQUARNAalt top-1 | 0.812 | 0.816 | 0.81 | 0.816 | 0.823 | 0.81 | 0.808 | 0.81 | 0.81 | 7.7 |
|  | SQUARNAsk top-1 | 0.815 | 0.827 | 0.809 | 0.814 | 0.824 | 0.804 | 0.816 | 0.831 | 0.814 | 9.9 |
|  | SQUARNA top-5 | 0.855 | 0.871 | <u>0.847</u> | 0.859 | 0.872 | <u>0.847</u> | 0.852 | 0.871 | <u>0.848</u> | 10 |
|  | **SQUARNAalt top-5** | 0.842 | 0.858 | 0.832 | 0.844 | 0.858 | 0.831 | 0.84 | 0.859 | 0.834 | 8.1 |
|  | **SQUARNAsk top-5** | <u>**0.857**</u> | <u>**0.879**</u> | 0.842 | <u>**0.862**</u> | <u>**0.883**</u> | 0.841 | <u>**0.853**</u> | <u>**0.876**</u> | 0.844 | 9.2 |
| **sequence**<br>**+ 2A3** | **RNAfold** | 0.633 | 0.71 | 0.587 | 0.666 | 0.72 | 0.619 | 0.603 | 0.701 | 0.558 | 0.6 |
|  | **RNAsubopt top-5** | 0.646 | 0.719 | 0.601 | 0.678 | 0.731 | 0.632 | 0.617 | 0.708 | 0.572 | 0.4 |
|  | **ShapeKnots top-1** | 0.745 | 0.751 | 0.744 | 0.767 | 0.772 | 0.762 | 0.724 | 0.732 | 0.726 | 1835.2 |
|  | **ShapeKnots top-5** | 0.817 | 0.827 | 0.815 | 0.836 | <u>**0.842**</u> | 0.831 | 0.799 | 0.812 | 0.799 | 1820.4 |
|  | **SQUARNA top-1** | 0.772 | 0.75 | 0.799 | 0.773 | 0.751 | 0.796 | 0.772 | 0.749 | 0.802 | 10.6 |
|  | **SQUARNAalt top-1** | 0.841 | 0.824 | 0.863 | 0.834 | 0.81 | 0.86 | 0.849 | 0.839 | 0.867 | 7.9 |
|  | **SQUARNAsk top-1** | 0.75 | 0.726 | 0.778 | 0.751 | 0.729 | 0.775 | 0.749 | 0.724 | 0.781 | 10.1 |
|  | **SQUARNA top-5** | 0.818 | 0.825 | 0.818 | 0.816 | 0.817 | 0.815 | 0.821 | 0.833 | 0.822 | 10.9 |
|  | **SQUARNAalt top-5** | <u>**0.845**</u> | 0.829 | <u>**0.865**</u> | <u>**0.838**</u> | 0.815 | <u>**0.862**</u> | <u>**0.853**</u> | 0.844 | <u>**0.869**</u> | 8.2 |
|  | **SQUARNAsk top-5** | 0.825 | <u>**0.838**</u> | 0.818 | 0.824 | 0.832 | 0.815 | 0.826 | <u>**0.845**</u> | 0.822 | 10.4 |

\*SPOT-RNA has a ~20 sec overhead time for a single run. We run the predictions sequence by sequence, resulting in the occurrence of overhead at each run. If all sequences were processed in a single bulk run, the runtime would be close to the time required for the prediction of the longest sequence (shown in parentheses)

## Notes

https://github.com/febos/SQUARNA

### Competing Interest Statement

The authors have declared no competing interest.

### Summary of Updates

Added an additional benchmark in terms of significantly covarying base pairs (Table S7, Figure S4)

